# Universality of representation in biological and artificial neural networks

**DOI:** 10.1101/2024.12.26.629294

**Authors:** Eghbal Hosseini, Colton Casto, Noga Zaslavsky, Colin Conwell, Mark Richardson, Evelina Fedorenko

## Abstract

Many artificial neural networks (ANNs) trained with ecologically plausible objectives on naturalistic data align with behavior and neural representations in biological systems. Here, we show that this alignment is a consequence of convergence onto the same representations by high-performing ANNs and by brains. We developed a method to identify stimuli that systematically vary the degree of inter-model representation agreement. Across language and vision, we then showed that stimuli from high-and low-agreement sets predictably modulated model-to-brain alignment. We also examined which stimulus features distinguish high-from low-agreement sentences and images. Our results establish representation universality as a core component in the model-to-brain alignment and provide a new approach for using ANNs to uncover the structure of biological representations and computations.

## Introduction

*“All happy families are alike; each unhappy family is unhappy in its own way.” Anna Karenina, Leo Tolstoy, 1877*

Across the tree of life, species have evolved convergent computational and behavioral strategies for interacting with their environment (Clark & Demb, 2016; Niven & Chittka, 2016). For example, humans and rainforest katydid insects, despite their distant phylogenetic histories, share remarkable similarity in the anatomy and function of the system for transforming air vibrations into a neural signal for hearing (Hoy, 2012; Montealegre-Z *et al*., 2012). Against the backdrop of numerous instances of biological convergence, artificial neural networks (ANNs) emerged in recent years, which mirror many behaviors observed in biological systems, including ones long been argued to be uniquely human, such as language (Devlin et al., 2018; LeCun et al., 2015; Mahowald et al., 2024; Radford et al., 2019). Despite differences in the hardware, learning algorithms, and resource constraints, in the presence of similar inputs and task demands, these ANNs appear to construct similar computational machinery to that of biological brains (Banino et al., 2018; Kell et al., 2018; P. Y. Wang et al., 2021). Indeed, numerous studies have found similarity in representations and behavior in direct comparisons between artificial and biological networks across perception, action, and cognition (Kell et al., 2018; Schrimpf et al., 2021; Sussillo et al., 2015; P. Y. Wang et al., 2021; Yamins et al., 2014). The fact that many different ANNs exhibit representations similar to the brain raises an intriguing possibility: that ANNs and brains are *converging onto universal representational axes* in the relevant domain. If true, this convergence would have significant implications for our understanding of both artificial networks (Dravid et al., 2023; Goh et al., 2021; Gurnee et al., 2024; Olah et al., 2020) and biological brains (Lindsay & Bau, 2023).

Previous studies comparing artificial and biological networks have left the question of representation universality open. Many studies have focused on a particular neural network model, designed to match the training data or the hypothesized objective to the brain, and evaluated the degree of similarity between them (Goldstein et al., 2022; Kell et al., 2018; Mano et al., 2021; Richards et al., 2019; Turner et al., 2019); for a review, see (Yamins & DiCarlo, 2016). Some studies have considered multiple networks and found that across domains, many high-performing ANN models capture biological representations comparably well (Caucheteux & King, 2022; Conwell, Prince, et al., 2024; Schrimpf et al., 2021, 2018; Tuckute et al., 2024; Yamins et al., 2014; Zhuang et al., 2021). Such findings could suggest that different networks are converging onto core domain-relevant representational axes (the ***representation universality*** hypothesis). However, they are also compatible with the possibility that the representations differ across models (the ***representation individuality*** hypothesis), but the available data fail to highlight these differences. Indeed, studies that have used carefully designed stimuli have revealed that ANNs do not always align with humans (Feather et al., 2023; Golan et al., 2020). To distinguish between these hypotheses, we developed a method to find stimuli that control the degree of representation agreement across ANNs and then used fMRI recordings to evaluate how well different models capture neural responses to these stimuli.

First, in the domain of language, we examined brain responses to stimuli (sentences) that elicited maximally divergent (i.e., individualized/model-specific) representations in several high-performing ANNs to test whether one particular ANN would come out as the best model of the human language system. We found instead that none of the ANN models could accurately predict brain responses to these low-agreement stimuli. We then tested whether responses to high-agreement stimuli, whose representations are convergent (i.e., universal) across models, would be better predicted. We evaluated this question for language stimuli, but also leveraged an existing dataset (Allen *et al*., 2022) to extend this approach to the vision domain. For both language and vision, we found that individual models predict brain responses more accurately for high-agreement, representationally-convergent stimuli than for low-agreement, representationally-divergent stimuli. Moreover, these findings hold in the other direction: stimuli that are represented similarly across individual brains are easier for models to predict. Finally, we took initial steps to identify the features that characterize representationally-convergent stimuli in language and vision.

Together, our findings provide evidence for ***representation universality*** among ANNs, and between artificial and biological networks, despite the stark differences in the underlying architecture, learning algorithms, and resource constraints. These shared representational axes are presumably determined by the input structure in the relevant domain (language, vision, etc.). The general approach that we developed for distilling the core representational axes across diverse state-of-the-art ANNs promises new insights into the nature of representations in biological systems.

## Results

### A novel method for identifying stimuli that differentiate ANNs

First, in the domain of language, we tested the hypothesis that one particular ANN model would significantly outperform other models in capturing brain responses to stimuli that are represented in distinct ways across models (the *representation individuality* hypothesis, **Figure 1A**). This outcome is expected if different models align with human neural representations for different reasons. In particular, only one model uses the same representational axes (feature space) as humans, but in most stimuli, the relevant features co-vary with some other, irrelevant features, which are the features that are captured by other models, leading to the result that all models can predict human responses well. To make this concrete, suppose one axis for human linguistic representations has to do with meaning abstractness, but it so happens that sentences expressing abstract meanings tend to be, on average, longer. In this scenario, one model represents the sentences in terms of the relevant, meaning-abstractness axis (similar to humans), but other models instead pick up on the irrelevant, sentence length, feature.

**Figure 1.**
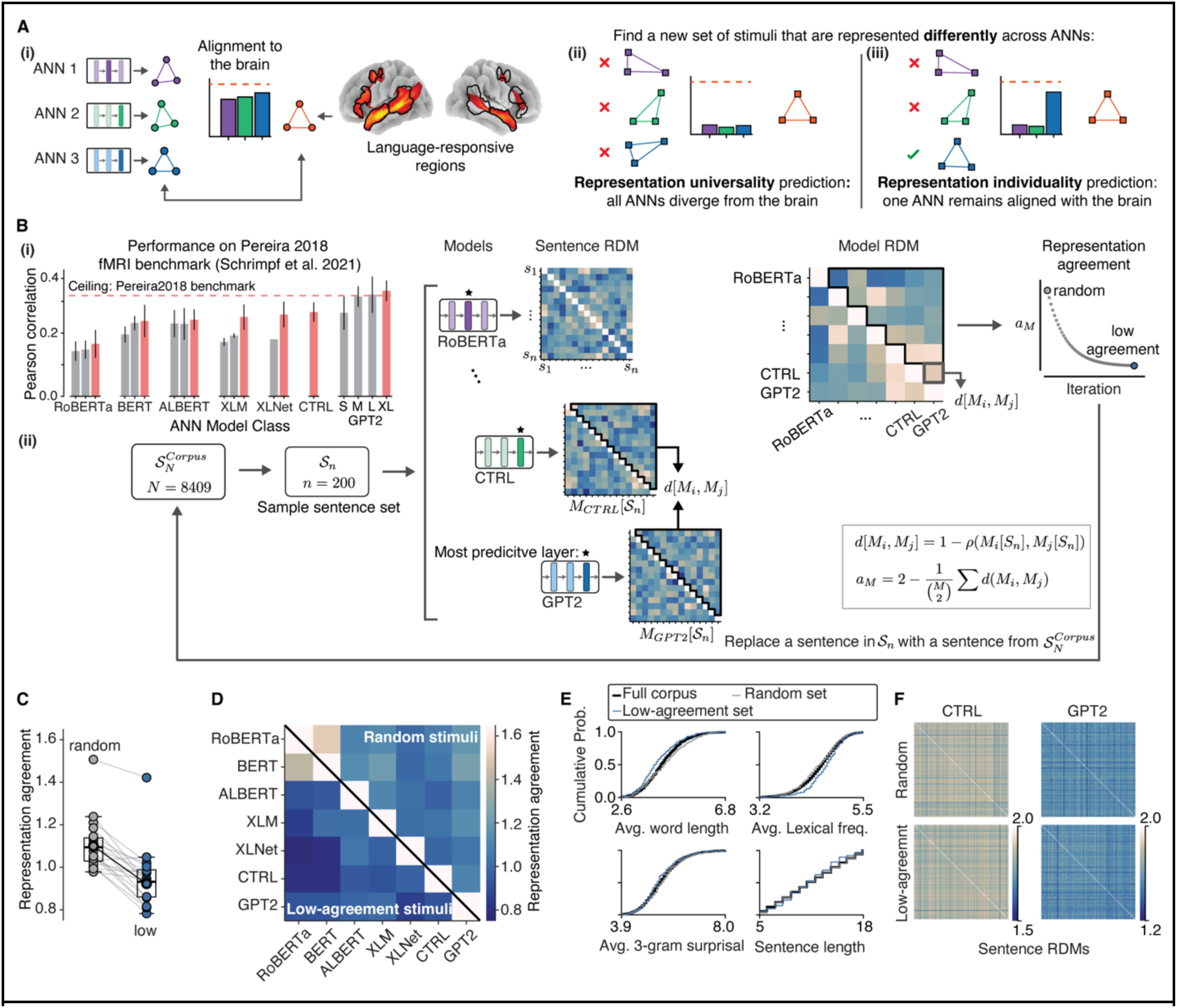
**A.** Representation universality and representation individuality as alternative hypotheses about why multiple ANNs align with brain representations for commonly used stimuli, which may not allow for model discrimination. Prior work has shown that many ANNs align with brain representations to a similar degree (A-i; language-responsive regions are shown because our focus is on language in Experiment 1), but why this happens is not clear: is it that all these ANNs are equally good models of the system in question, or is it that we are failing to discriminate among them? Distinguishing between these hypotheses requires stimuli that are represented differently across models. For these critical stimuli, under the representation universality hypothesis, no model should align with brain data given that inter-model divergence should also lead to divergence from the brain (A-ii). Alternatively, under the representation individuality hypothesis, one model should align with the brain regardless of its low agreement with other models (A-iii). **B.** A method for selecting stimuli that are represented differently across models. (B-i) Performance of seven classes of ANN models (measured with cross-validated Pearson correlation; <u>Methods</u>) on the *Pereira2018* fMRI benchmark, as reported in (Schrimpf *et al*., 2021). The highest-performing model variants (in red) were chosen for stimulus optimization. (B-ii) Stimulus sampling approach. Briefly, a set of **n** sentences were sampled from a corpus and the Sentence RDM was computed for each model, using the layer that best predicted fMRI activation in Schrimpf et al. (2021). The upper triangular parts of the Sentence RDMs were then compared across models using Pearson correlation to construct a Model RDM. The Model RDM quantifies the degree of agreement for every model pair. The average of the upper triangular part of the Model RDM was then minimized (or maximized, for high-agreement stimuli) by iteratively swapping sentences with replacements from the corpus. **C.** Comparison of representation agreement across model-pairs (7 models, 21 model pairs) for an average random set of sentences and the low-agreement set. We constructed an average random set by sampling 200 sets of random sentences (200 sentences each) and measuring model-pair agreements; we then averaged each model-pair agreement across the 200 samples to arrive at an average model-pair agreement. For most model pairs, the agreement was lower (darker blue colors) as a result of the stimulus optimization procedure. The dark solid line shows the change in agreement for CTRL and GPT2 model pairs. **D.** Individual model-pair representation agreements for the random (upper triangular) and low-agreement sentence sets (lower triangular). **E.** Empirical cumulative probability distributions for several linguistic features in the full corpus, a sample random set of sentences, and the low-agreement set. No significant differences were observed between the low-agreement and random sets, except for average lexical frequency (<u>Methods</u>, see **Suppl. Figure 1B** for additional linguistic features). **F.** The Sentence RDMs of two example models (CTRL and GPT2) for random and the low-agreement sentences. The distributions of Sentence RDM values in each model were similar between the random and low-agreement sentence sets (except for BERT and RoBERTa; RDMs for all models are shown in **Suppl. Figure 1A**).

According to the *representation individuality* hypothesis then, only the former model will be able to predict neural responses to stimuli that dissociate these two axes. On the other hand, according to the *representation universality* hypothesis, all ANNs should struggle to predict brain responses to such stimuli because any model-specific representation would be distinct not only from the representations of other models but also from the brain representations (**Figure 1A**).

For testing ANNs against brain responses, we adopted a critical criterion from the strong inference approach (Platt, 1964) whereby each model provides a distinct hypothesis about how linguistic input is represented. To this end, we devised an approach for stimulus selection based on representational dissimilarity. We additionally constrained the selection method to only span ecological, or naturally occurring, stimuli (in this case, sentences—the preferred stimulus of the human language network; (Fedorenko et al., 2024) because we wanted to ensure that we would be able to obtain strong and reliable brain responses, which may be more difficult for artificially constructed stimuli (e.g., (Golan et al., 2020, 2023).

We started by extracting internal representations from a cohort of language ANNs (large language models) for a set of sentences. Following prior work (e.g., Schrimpf et al., 2021), we chose seven architecturally distinct models that provided the best in-class predictivity of fMRI voxel-level responses in the language network (RoBERTa: roberta-base (Liu *et al*., 2019), BERT: bert-large-uncased-whole-word-masking (Devlin *et al*., 2018), ALBERT: albert-xxlarge-v2 (Lan *et al*., 2019), XLM: xlm-mlm-en-2048 (Lample & Conneau, 2019), XLNET: xlnet-large-cased (Yang *et al*., 2019), CTRL: ctrl (Keskar *et al*., 2019), GPT2: gpt2-xl (Radford *et al*., 2019); **Figure 1B**). Aside from their architecture, these models vary in their training objective, training dataset, and tokenization of inputs (<u>Methods</u>). We then used representational dissimilarity (Kriegeskorte & Diedrichsen, 2019; Kriegeskorte et al., 2008; Sucholutsky et al., 2023) to select sentences that would be represented as differently as possible across these seven models. We chose representational dissimilarity because i) language ANNs have different numbers of dimensions to represent the input, which makes direct comparisons of embeddings challenging; and ii) comparing two models in terms of their representational dissimilarity is symmetric, unlike regression-based approaches (Harvey et al., 2023; Williams et al., 2021).

Because representational dissimilarity is quantified across multiple stimuli, we defined a representation agreement measure for a *set* of sentences. In particular, for each model, we computed pairwise sentence dissimilarities for 200 sentences using a correlation coefficient (<u>Methods</u>). We then defined agreement between two models using the correlation between their sentence representational dissimilarity matrices (RDMs). We computed agreement for every pair of models (21 pairs) and subsequently optimized the average of inter-model representation agreements.

To identify sentences for which model representations maximally diverge, we started with a random set of 200 sentences from a corpus of 8,409 naturally occurring sentences (the Universal Dependencies corpus (de Marneffe et al., 2021); <u>Methods</u>). We then iteratively substituted a sentence from the remaining sentences and updated the set if the substitution decreased representation agreement across ANNs. This process was repeated until we reached the minimum agreement value for this corpus (**Figure 1B**). We call the resulting sentences “low-agreement sentences”.

For these low-agreement sentences, the pairwise model similarity was lower compared to an average random set of 200 sentences from the same corpus (**Figure 1C,D**), and to sentences used in prior neural benchmarks, indicating that the optimization procedure was successful (**Figure 2A**; <u>Methods</u>) (low-agreement set median pairwise model agreement: 0.931; random set: 1.094; *Pereira2018* benchmark (from (Schrimpf et al., 2021): 1.124 and 1.136 for the two sentence sets, Mann-Whitney U-tests for the differences between the low-agreement set and each of the other sets: *p*s<0.001).

**Figure 2.**
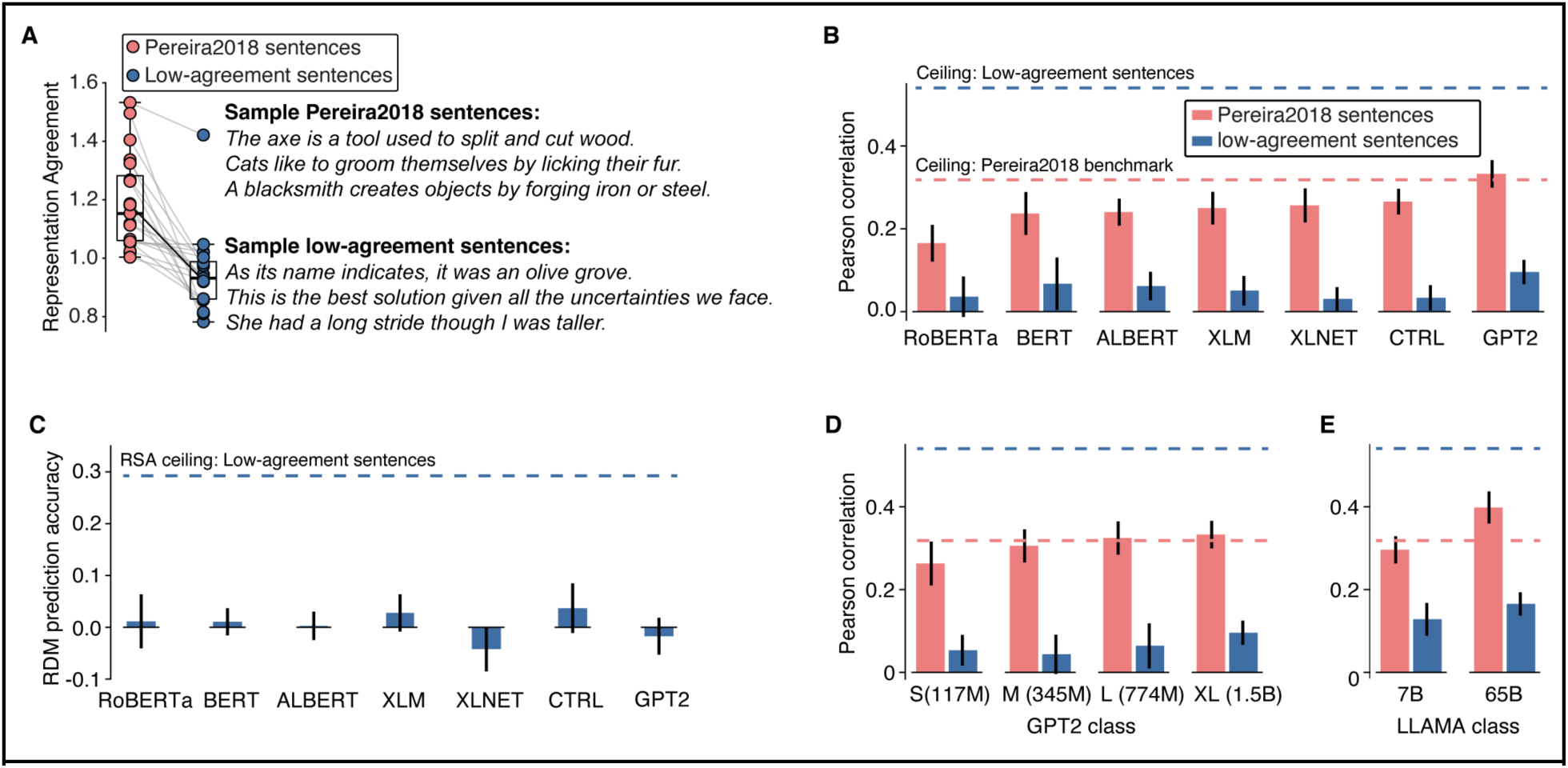
**A.** Representation agreement across model pairs (individual lines) for the sentences used in the *Pereira2018* fMRI benchmark and the low-agreement sentences. The level of agreement was substantially higher for the *Pereira2018* sentences. The dark solid line shows the change in agreement for CTRL and GPT2 model pairs. **B.** Performance of ANN models in predicting voxel responses in language-responsive regions for the *Pereira2018* sentences (red; (Schrimpf et al., 2021) and the low-agreement sentences (blue), as estimated with an encoding model. Ceilings based on inter-subject correlations (<u>Methods</u>) for each set are shown with dashed lines. Consistent with the *representation universality* hypothesis, all models struggle to predict voxel responses for the low-agreement sentences, as evidenced by low correlations and a much larger gap between the correlations and the ceiling compared to the *Pereira2018* benchmark. (Note that for the *Pereira2018* benchmark, predictivity is close to or above the ceiling level for some models—this is likely because performance was overestimated in the original paper due to the cross-validation procedure used (see (Kauf *et al*., 2023). **C.** Performance of ANN models in predicting RDMs constructed from voxel responses across the language-responsive regions. All models struggle to predict voxel RDMs, with performance dropping to baseline, compared to the ceiling at 0.3 (based on leave-one-out RDM similarity across participants, <u>Methods</u>). **D-E.** Generalization of the results to models that were not used for stimulus selection, including models in the GPT2 class (S, M, and L, with the XL model shown for comparison) (D) and newer, large-scale models (LLAMA models with 7 billion and 65 billion parameters; compared to GPT2-XL, which has 1.5 billion parameters) (E). All models struggle to predict voxel responses for the low-agreement sentences.

No salient qualitative differences are apparent between the low-agreement and random sentences: sentences in the low-agreement set, similar to a random sample, are grammatically well-formed, use diverse structures, and span different meanings. To quantitively evaluate whether the low-agreement sentences fell outside the distribution of natural language, we evaluated them using standard linguistic metrics (average word length, lexical frequency and n-gram surprisal, sentence length in words, etc.) and found that they were highly overlapping with the full corpus and with 200 subsets of random sentences (**Figure 1E**, see **Suppl. Figure 1B** for other linguistic features). To quantify this overlap, we performed Mann-Whitney U-tests, for each feature, comparing the low-agreement set against each of the 200 random subsets (<u>Methods</u>). The proportion of tests showing significant differences provided empirical p-values. For most linguistic features, the distributions in the random sets were similar to the distribution for the low-agreement set (*p*s>0.09; except average lexical frequency, which showed a small but significant effect, **Suppl Figure 1B**). Finally, to evaluate whether the low-agreement sentences were outliers in the models’ representational spaces, we compared their representational geometry, in the form of RDMs, to that of random sentences using the same approach as above (Methods). The RDM distributions were similar for 5 of the 7 models (*p*s>0.2, except for RoBERTa: p=0.045, and BERT: p=0.01).

### Representational divergence among ANNs leads to divergence from brain representations for language

In Experiment 1, we collected fMRI responses to the 200 low-agreement sentences from 8 participants (one 2-hour scanning session each, two presentations of each sentence, **Suppl. Figure 1C**). For the analyses, we focused on the language-responsive regions identified in each participant with an independent localizer task (Fedorenko et al., 2010; <u>Methods</u>). For the voxels in the language areas, we measured the reliability of responses by computing a correlation (across the 200 sentences) between the two repetitions, and selected voxels in the top 10% of the correlation values, excluding the small number of voxels for which the correlation did not significantly differ from a shuffled baseline (<u>Methods</u>). This selection procedure left 1,093 voxels across participants (average=137 voxels per participant), with the median correlation value of r=0.27 (SD=0.13) (**Suppl. Figure 1D**, <u>Methods</u>). These voxels showed a strong average BOLD response to the 200 critical sentences, as expected given the strong response in this system to language stimuli (first repetition: median = 0.39 beta values, SD = 4.7; second repetition: median = 0.35, SD = 4.8), whereas the response was notably lower for the 16 nonword sequences interspersed with sentences (first repetition: median =-0.21 beta values, SD = 4.5; second repetition: median = 0.04, SD = 4.5; Mann-Whitney U-tests for the differences between sentences and nonwords: p-value < 0.001). We also measured the consistency of neural responses across participants (<u>Methods</u>) and found high inter-individual consistency, similar to prior work (Pearson correlation of 0.54 between BOLD responses of a given participant and predicted responses based on the remaining participants, reported as ceiling in (Schrimpf *et al*., 2021; Tuckute *et al*., 2024).

Next and critically, we tested the ANN models’ ability to predict neural responses and found that no model could predict responses to the low-agreement sentences (**Figure 2B**). To evaluate model-to-brain similarity, we used the standard encoding model approach: in each participant, we constructed a 5-fold cross-validated linear regression between model activations and voxel responses in individual participants (Schrimpf *et al*., 2021), and computed a Pearson correlation between predicted and observed responses for each fold. The final Pearson correlation values—averaged across folds, voxels, and then participants—were low for all models, as evidenced by the large gap between each model’s predictivity and the ceiling for the low-agreement set; they were also substantially lower than previously reported results for the *Pereira2018* benchmark of fMRI responses to semantically diverse sentences (Schrimpf *et al*., 2021) (**Figure 2B**, see **Suppl. Figure 1G-J** for predictivity across all layers of representative models; see **Suppl. Figure 2E** for evidence that the discrepancy is not due to visual vs. auditory presentation modality). In other words, no individual model excelled at predicting brain responses for these representationally-divergent stimuli. The results generalize to a different model-to-brain comparison approach: representational similarity analysis (RSA; (Kriegeskorte et al., 2008; Schütt et al., 2023). RSA correlations were close to zero for all models (**Figure 2C**), well below the inter-individual consistency measured by RSA (reported as the RSA ceiling, <u>Methods</u>).

The results so far provide evidence against the possibility that one of the seven models is a better model of the language system (the *representation individuality* hypothesis) and are instead consistent with—though do not yet directly test—the *representation universality* hypothesis. But perhaps our set of high-performing ANNs simply doesn’t include the right model of the language system. Although this possibility is difficult to rule out completely, we next tested whether the low model-to-brain similarity results would generalize to models outside of the original set. Under the *representation universality* hypothesis, all high-performing ANNs converge onto the same representational axes, so we would expect that even for ANNs outside of our initial set, low-agreement stimuli would still produce poor model to brain-alignment. First, we considered models from the same class as GPT2-XL that were not used for stimulus selection (i.e., in selecting the sentences whose representations are maximally divergent across models). We found that 3 additional variants of GPT2 that differ in their number of parameters (GPT2-S, GPT2-M, GPT2-L) similarly fail to predict brain responses to the low-agreement sentences (**Figure 2D**). Importantly, these models predicted brain responses to the *Pereira2018* sentences on par with GPT2-XL (**Figure 2D**, (Schrimpf *et al*., 2021). Second, we considered a model class outside of the original set and with substantially more model parameters and training: LLAMA-7B and LLAMA-65B that have 5- and 50-fold more parameters, respectively, than the largest model in our set (GPT2-XL, 1.5 billion parameters) and were trained on up to 1.4 trillion tokens (Touvron *et al*., 2023). For both models, the ability to predict responses to the low-agreement sentences was far below that for the *Pereira2018* benchmark (**Figure 2E**). These results strengthen our confidence that no individual ANN model can predict brain responses for the low-agreement sentences.

### Representational convergence among ANNs leads to convergence with brain representations for both language and vision

We have so far demonstrated that when models are in low agreement, no individual ANN model emerges as the best model of the human language network: all models fail to predict the language areas’ responses to the low-agreement sentences. These results are compatible with the *representational universality* hypothesis, but they do not yet directly test it. An alternative explanation for the results could be that the representation agreement metric modulates features across ANNs that don’t contribute to model-to-brain alignment (**Figure 3A)**. To evaluate the *representational universality* hypothesis more explicitly, we next asked whether we can modulate model-to-brain alignment in the other direction: namely, *increase* it by examining brain responses to sentences whose representations are similar across ANNs (i.e., universal). This positive result would establish that the representational axes are shared between models and brains, such that by modulating the level of inter-model representation agreement we can decrease or increase the level of model-to-brain alignment.

**Figure 3.**
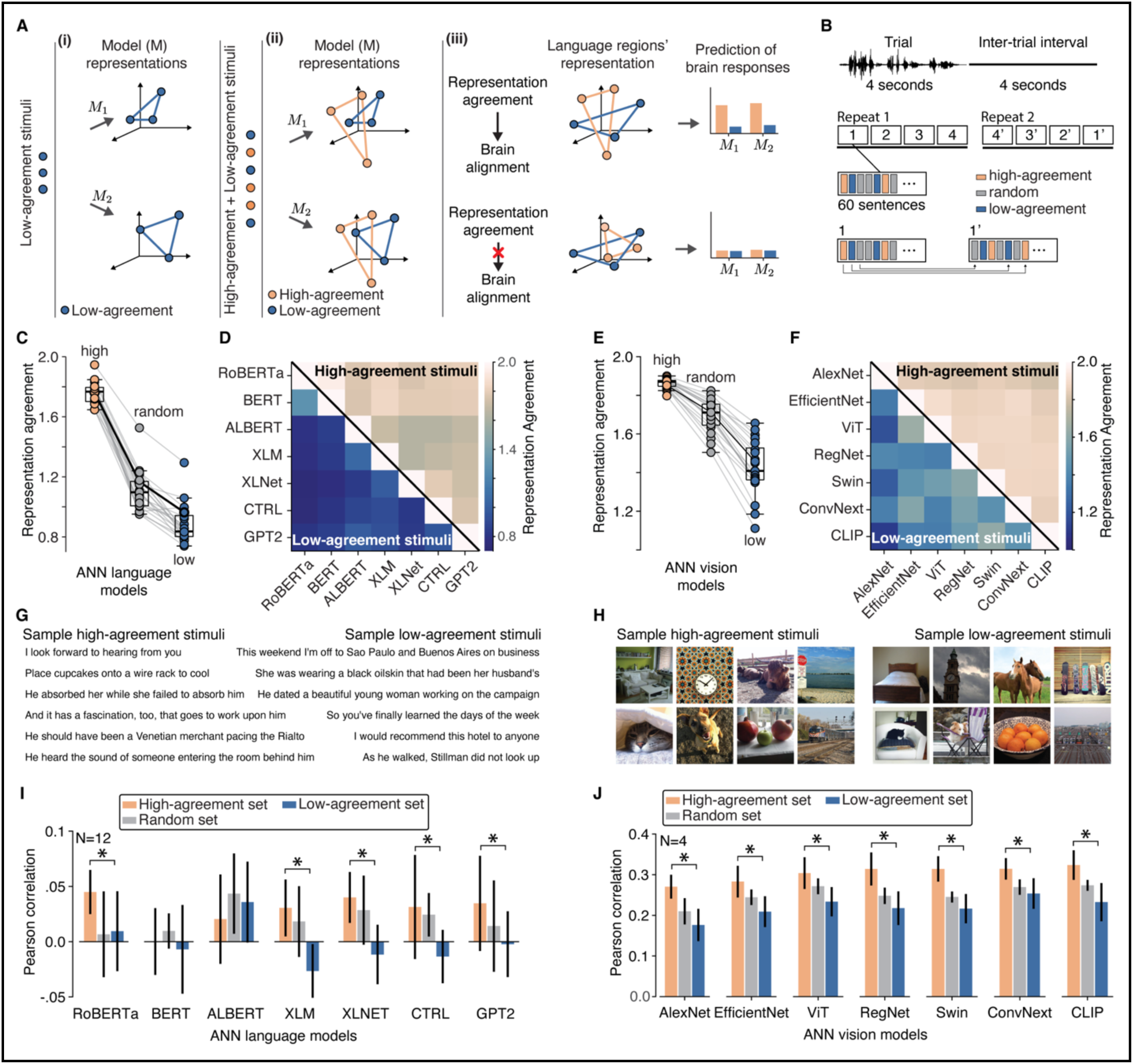
**A.** Two possible relationships between inter-model representation agreement and model-to-brain alignment. (A-i) Hypothetical representations for three low-agreement stimuli in two models (M1 and M2). (A-ii) Hypothetical representations for three high-agreement (orange) and three low-agreement (blue) stimuli in two models. The geometry of the high-agreement stimuli is consistent across models (the orange triangles have similar shapes). The geometry of the low-agreement stimuli is not consistent across models (the blue triangles have different shapes). (Note that this visualization is not meant to imply that the two groups of stimuli reside in distinct subspaces of model representation; we are simply highlighting their different configurations in the representational space.) (A-iii) If inter-model representation agreement modulates model-to-brain alignment (i.e., individual models are better at predicting voxel responses to the high-agreement than the low-agreement stimuli; Top), that would suggest that the inter-model representation agreement measure is successfully tapping the shared aspects of model and brain representations. Otherwise (Bottom), the inter-model representation agreement measure is not a good approach for tapping the shared aspects of model and brain representations. **B.** Experimental design. Each trial consisted of a 4 second audio of a sentence, and a 4 second inter-trial interval period. The scanning session consisted of 2 repetitions of each stimulus, distributed over 8 runs (60 sentences each: a combination of high-agreement, random, and low-agreement sentences presented in pseudo-random order). The order of sentences varied between repetitions (1 and 1’). **C.** Representation agreement across model pairs (individual lines) for the high-agreement, random, and low-agreement sentences. The dark solid line shows the change in agreement for CTRL and GPT2 model pairs. **D.** Individual model-pair representation agreements for the high-agreement (upper triangular) and low-agreement sentence sets (lower triangular). **E-F.** Same as (C-D) but for the visual stimuli. The dark solid line shows the change in agreement for ConvNext and CLIP model pairs. **G-H.** Example high-agreement and low-agreement sentences (G) and images (H). **I.** Performance of ANN language models in predicting voxel responses in language-responsive regions for the high-agreement sentences (orange) and low-agreement sentences (blue; performance for the random set is shown in grey). Consistent with the *representation universality* hypothesis, for 5 models, the Pearson correlation is significantly higher than for the high-agreement than the low-agreement set, with the random set falling in between. **J.** Performance of ANN vision models in predicting voxel responses in high-level visual regions for the high-agreement images (orange) and low-agreement images (blue; performance for the random set is shown in grey). Consistent with the *representation universality* hypothesis, for all models, the Pearson correlation is significantly higher for the high-agreement than the low-agreement set, with the random set falling in between.

We adapted our stimulus selection approach (**Figure 1B**) to find a set of sentences for which model representations maximally converge across the same seven ANNs (<u>Methods</u>). In addition to these high-agreement sentences, we selected another set of low-agreement sentences (non-overlapping with the set in Experiment 1), and a set of random sentences (80 sentences in each set; see **Figure 3G** for sample sentences). The pairwise model similarity differed reliably among the three sets for all model pairs (**Figure 3C, D**), indicating that the optimization procedure was successful (high-agreement set median pairwise model agreement: 1.767; random set: 1.097; low-agreement set: 0.836; Mann-Whitney U-tests: *p*s<0.001). We evaluated the three sets of sentences using the same linguistic metrics as in Experiment 1 and found no reliable differences for the critical comparison between the high-and low-agreement sets, except for the average lexical frequency, which showed a small significant difference (**Suppl. Figure 2B**).

Similar to Experiment 1, we also tested for each language model whether the distribution of RDM values deviates from what would be expected under random sampling and found that it does not (empirical p-values > 0.075; Methods).

In Experiment 2, we collected fMRI responses to the three sets of sentences from 12 participants (one 2-hour session each, two presentations of each sentence). As in Experiment 1, we focused on the language areas, identified with a localizer task (Fedorenko *et al*., 2010; <u>Methods</u>), and restricted our analyses to voxels that show reliable responses across sentence repetitions (reliability was computed using only the random sentence set, to remove any bias toward voxels that might respond differentially across different sets; <u>Methods</u>). This selection procedure left 1,086 voxels across participants (average=91 voxels per participant), with the median correlation value of 0.24 (SD=0.052; **Suppl. Figure 2D**). As expected, these voxels showed a strong average BOLD response to the three conditions **(Suppl. Figure 2C)**. Finally, we measured the consistency of neural responses across participants, as in Experiment 1, for each condition separately (<u>Methods</u>) and found similar consistency values across the high-agreement, random, and low-agreement sets (high-agreement: 0.244; random: 0.206; low agreement: 0.249; <u>Methods</u>). To measure alignment between ANNs and the brain, we performed individual 5-fold cross-validated regression on first and second repetition of stimuli in each stimulus group independently and averaged the regression score across repetitions.

We found that for the majority of the models, the prediction of responses for the high-agreement sentences was substantially higher than for the low-agreement sentences (**Figure 3I**). For five of the seven models (RoBERTa, XLM, XLNET, CTRL, GPT2), this difference was significant (*p*s<0.05; **Figure 3I**; see **Suppl. Figure 2E** for the results for the individual participants; see **Suppl. Figure 2G,H** for evidence that the patterns are similar regardless of whether the model predictions are normalized by across-participant ceilings, including when ceilings are computed separately for each condition). The results generalize to ANNs that were not used for stimulus selection (e.g., GPT-2 medium and large; **Suppl. Figure 2F**). Moreover, the relationship between representational agreement and predictivity of brain responses is only observed for high-performing ANN language models. In particular, the difference in predictivity between the high-agreement and low-agreement sentences is not observed in several baseline models (a random embedding, a decontextualized embedding (word2vec; Mikolov, et al., 2013), and an LSTM model; (Jozefowicz *et al*., 2016), which all showed poor ability to predict fMRI brain responses for the *Pereira2018* benchmark in Schrimpf et al. (2021) (**Suppl. Figure 2I**). In addition, for transformer models, the difference only emerges in larger models, but not in GPT-S or distilGPT2 (**Suppl. Figure 2F**), which suggests that convergence onto the same representational axes only characterizes high-performing ANN language models. Finally, the results were spatially restricted to the language brain areas, as revealed by a control analysis of lower-level auditory areas, which respond strongly to auditorily presented language stimuli but do not show the same pattern of model predictions (**Suppl. Figure 2K**).

To test the broader generality of the *representational universality* hypothesis, we turned to a different domain: vision. Similar to how multiple high-performing language ANNs can predict neural responses in the language brain areas (Schrimpf *et al*., 2021), many neural network vision models can predict neural responses in the ventral visual cortex (Conwell, Prince, et al., 2024) Following our approach above, we selected seven ANN vision models from diverse architectural classes that provided the best in-class predictivity of fMRI voxel-level responses in human occipitotemporal cortex (OTC) (Allen et al., 2022; Conwell, Prince, et al., 2024). We modified the stimulus selection algorithm with an additional criterion to keep the distribution of image dissimilarities within each ANN close to what would be expected from a random sample of images (using KL divergence as a penalty term; <u>Methods</u>). We then selected three sets of 80 images from the set of 1,000 images used in (Allen *et al*., 2022): the low-agreement image set, a random set of images, and the high-agreement image set (**Figure 3E, F**). No salient qualitative differences are apparent among the sets: all sets contain diverse images spanning different categories (faces, natural scenes, food, etc.; **Figure 3H**). Similar to Experiment 2, we tested for each vision model whether the distribution of RDM values deviates from what would be expected under random sampling and found that it does not (empirical p-values > 0.29; see <u>Methods</u>).

In Experiment 3, we used fMRI data from (Allen *et al*., 2022) from 4 participants (43 scanning sessions each, resulting in three presentations of each image for the 1,000 images that were presented to all participants). We used a broad mask for the occipitotemporal cortex (also referred to as human IT), which includes all the well-characterized category-selective areas (Kanwisher, 2010), and restricted our analyses to voxels that showed response reliability of 0.2 or higher (reliability was computed as in (Allen et al., 2022; Conwell, Prince, et al., 2024; Tarhan & Konkle, 2020a). This selection procedure left 29480 voxels across participants (average=7370 voxels per participant; <u>Methods</u>). To evaluate model-to-brain similarity, we computed a Pearson correlation using a cross-validated regularized regression, to remain consistent with (Conwell, Prince, et al., 2024) (<u>Methods</u>).

Similar to what we found for the language domain, the prediction of responses for the high-agreement images was substantially higher than for the low-agreement images (**Figure 3J**). This difference was significant for all models (*p*s<0.05; **Figure 3J**). The results were spatially restricted to the higher-order visual brain areas, as revealed by a control analysis of lower-level visual areas, which responded strongly to visual stimuli but do not show the same pattern of model predictions (**Suppl. Figure 3B**). To evaluate the robustness of the results to how the mapping model is trained, we trained the mapping model on a much larger set of stimuli (n=774 mages, which were not used in any of the three sets), which, we reasoned, may better capture the universal parts of representational spaces in each model. Using this mapping model to predict the responses to high-and low-agreement stimuli revealed the same pattern of results: the prediction for the high-agreement images was higher than for the low-agreement images (**Suppl. Figure 3C**), and again, this effect was restricted to the higher-level visual areas (**Suppl. Figure 3D**).

To summarize so far, for both language and vision, ANN models can more accurately predict brain responses for high-agreement (representationally-convergent) stimuli compared to low-agreement (representationally-divergent) stimuli. One possible alternative explanation for our results is that brain responses to the stimuli (sentences or images) in the high-agreement sets might be easier to predict simply because the stimuli themselves are more similar to one another, thereby facilitating generalization from the training to the test stimuli in the encoding model regression analyses. To evaluate this possibility, we examined the relationship between the degree of inter-stimulus similarity and the Pearson correlation values from the encoding models (language: **Suppl. Figure 2K**; vision: **Suppl. Figure 3D**) and evaluated whether the set type explained variance in the brain predictivity over and above inter-stimulus similarity (evaluated with nested linear mixed-effects models (LMEs) and the likelihood ratio test, <u>Methods</u>). Both in language and vision, we found that set type explained additional variance in brain predictivity beyond inter-stimulus similarity (language: *x^2^*=21.06, p=2.66e-05; vision: *x^2^*=85.16, p=3.22e-19).

### Representational convergence among individual brains leads to convergence with ANN representations

In the previous sections, we found evidence for the *representational universality* hypothesis, such that brain responses to linguistic and visual stimuli that are represented similarly across ANNs are better predicted across models, compared to responses to stimuli that models represent differently. We next tested a prediction of this hypothesis in the direction from biological to artificial networks. In particular, we asked whether the degree of representational agreement across individual brains would increase model-to-brain alignment. This result is not guaranteed given the findings so far: inter-brain representational similarity as measured by fMRI may vary along axes that only partially overlap with the representational axes used by ANNs.

In Experiment 4, we focused on the vision domain given that we have access to brain responses for a large set of images (Allen et al., 2022). We treated each participant as a model and used our stimulus selection approach (**Figure 1C**) to find sets of images for which the fMRI responses were in either high or low agreement (**Figure 4A**). Importantly, these image sets were largely non-overlapping with the ones selected based on high or low inter-model agreement in the previous section (high-agreement: 15% overlap, random: 7.5%, low-agreement: 12.5%). We then tested how similar ANN representations were for these image sets and how well ANNs could predict fMRI responses to each set.

**Figure 4.**
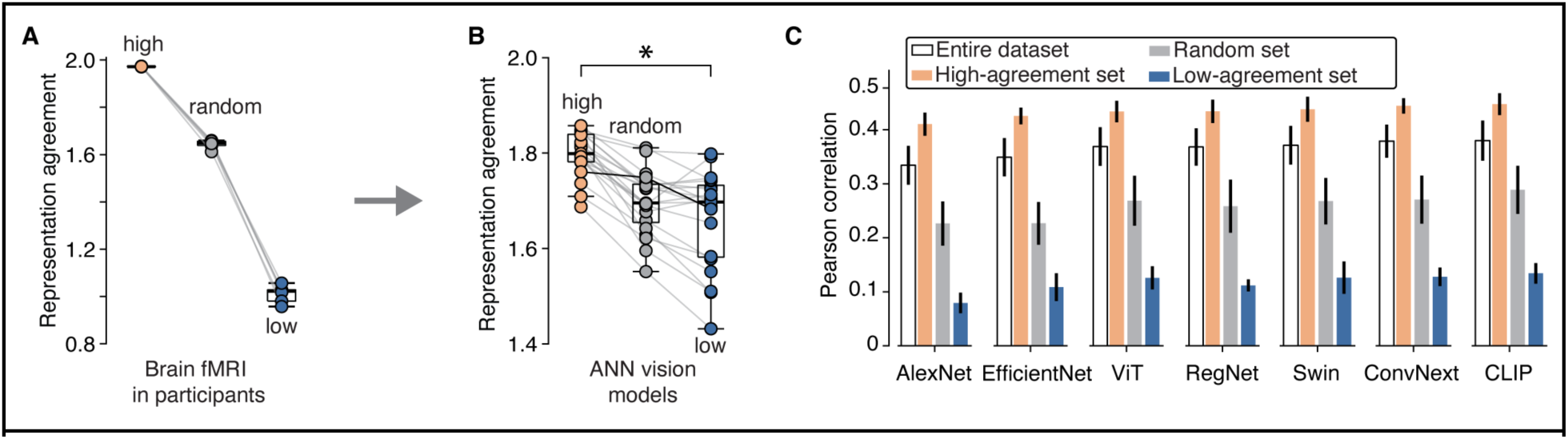
The relationship between representation universality across participants and models. **A.** Representation agreement in fMRI responses across participant pairs (individual lines; 4 participants, 6 participant pairs) for the high-agreement, random, and low-agreement images sets. **B.** Representation agreement across ANN model pairs (individual lines) for the high-agreement, random, and low-agreement images, which were selected based on inter-participant agreement in (A). The dark solid line shows the change in agreement for ConvNext and CLIP model pairs. **C.** Performance of ANN vision models in predicting voxel responses in high-level visual regions for the high-agreement image (orange) and low-agreement images (blue; performance for the random set is shown in grey), selected based on inter-participant agreement. Consistent with the *representation universality* hypothesis, for all models, the Pearson correlation is significantly higher for the high-agreement than the low-agreement set, with the random set falling in between.

We found that representation agreement across human participants indeed modulated representation agreement across the 7 ANN vision models (**Figure 4B**, medians: high-agreement=1.798, random= 1.695, low-agreement=1.697, Wilcoxon rank sum test *p*-value(high>low)<0.001, *p*-value(random>low)=0.0099). Moreover, all models predicted fMRI responses to the high-agreement images substantially better than to the random or low-agreement stimuli (**Figure 4C**). These results suggest that representational universality holds both from models to brains, and from brains to models.

### Dimensions of the universal representations in language and vision

We have so far presented evidence for representation universality by showing that stimuli that are convergent across ANN models align better with human brain representations, and that stimuli that are convergent across brains are also more similarly represented across ANNs.

What features distinguish high-agreement and low-agreement stimuli? By analyzing the properties of these different stimulus sets, can we infer something about the coding axes of linguistic and visual stimulus spaces? We undertook an initial exploration of this question using human behavioral ratings.

For language, prior to behavioral data collection, we examined model behavior across different sets, asking: how well can the models predict upcoming words at each position in the sentence. We found that for all models, next-word prediction performance was similar between the high-and low-agreement sentences (**Suppl. Figure 4A**). This result aligns with the fact that high-and low-agreement sentences show strong overlap in various linguistic features, such as average word frequency, examined in **Suppl. Figure 2B**, and rules out the possibility that the models are simply unable to process or represent the low-agreement sentences. We then collected human ratings for our three sentence sets (240 sentences total) (Rayner & Duffy, 1986) on a set of 10 stimulus features from n=360 participants recruited online (n=23-36 per feature). The properties were motivated by past work in psycholinguistics and brain imaging (Arfé et al., 2023; Binder et al., 2005; Demberg & Keller, 2008; Fedorenko et al., 2020; Ferstl & von Cramon, 2007; Kuchinke et al., 2005; Rayner & Duffy, 1986) and recent work showing that some of these features explain variability in the brain’s responses to sentences (Tuckute et al., 2024).They included both form-related features (e.g., grammaticality) and meaning-related features (e.g., whether the sentence talks about the physical world) (**Figure 5A**, <u>Methods</u>). One additional feature was included based on our intuitions about a possible difference between the high-and low-agreement sentences: what we termed ‘meaning generality’, or whether the sentence could be used in many different contexts (similar to the ‘contextual diversity’ measure for words; (Adelman et al., 2006) (**Figure 5A**).

**Figure 5.**
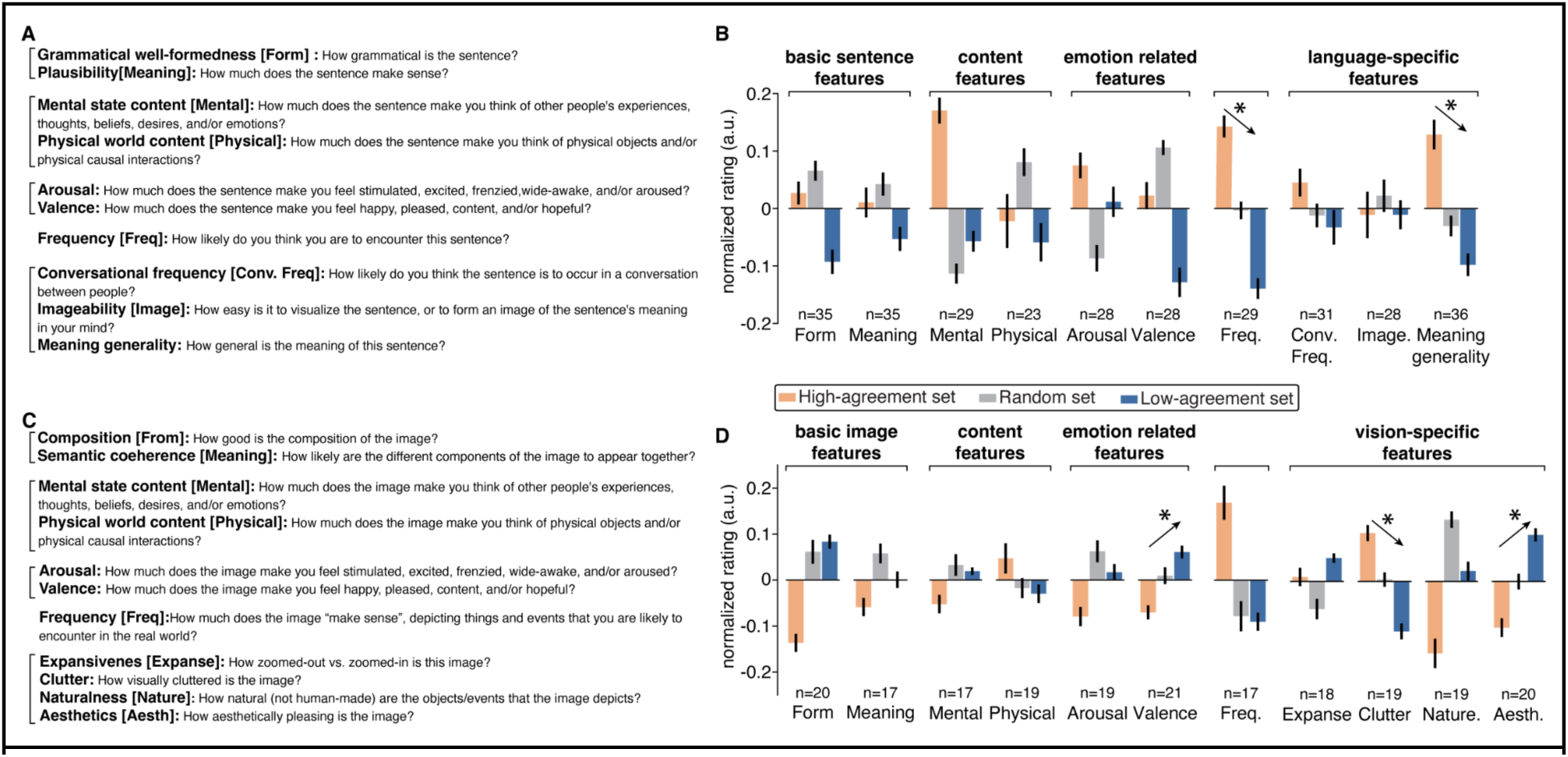
Searching for candidate features that vary between high-agreement and low-agreement stimuli. **A.** Human participants rated sentences on 10 dimensions using a Likert 1-7 scale. Each participant rated sentences on a single dimension, except for grammatical well-formedness and plausibility, where participants rated both (following Tuckute et al., 2024). **B.** Participant-normalized ratings of the three sentence sets on each dimension. For Perceived Frequency, the high-agreement sentences were rated as more frequent than the random sentences, which in turn, were rated as more frequent than the low-agreement sentences. Meaning generality showed a similar pattern between the high-, random-, and low-agreement sentences. Error bars show the standard error across participants. **C.** Human participants rated images on 11 dimensions using a Likert 1–7 scale. Each participant rated images on a single dimension. **D.** Participant-normalized ratings of the three image sets on each dimension. For Visual Clutter, the high-agreement images were rated as having more visual clutter than the random images, which in turn, were rated as having more visual clutter than the low-agreement images. For Valence and Aesthetics, the high-agreement images were rated as more negative and less visually appealing than the random images, which in turn, were rated as more negative and less appealing than the low-agreement images. Error bars show the standard error across participants.

For a feature to be interpreted as an axis that goes from the high-agreement part of the representation space to the low-agreement part, it would need to show a consistent direction of change with the change in the degree of representation agreement (e.g., feature(high-agreement sentences) > feature(random sentences) > feature(low-agreement sentences)) or the opposite trend (high-agreement < random < low-agreement)). For sentences, only two features exhibit this pattern (**Figure 5B**). One is *Perceived Frequency*: the high-agreement sentences were rated as more frequent than the random sentences, and the random sentences were, in turn, rated as more frequent than the low-agreement sentences (mean over participants: high-agreement: 0.14, random: ∼0.0, low-agreement:-0.14, *p*-value (high>random)<0.001, *p*-value (random>low)<0.001, paired-samples t-test) (*Perceived Conversational Frequency* showed a similar trend, but not reliably). The other feature is *Meaning Generality*: the high-agreement sentences were rated as higher in meaning generality than the random sentences, which in turn were rated higher than the low-agreement sentences (high-agreement: 0.13, random: ∼0, low-agreement:-0.09, *p*-value(high>random)<0.001, *p*-value(random>low)<0.05, paired-samples t-test).

For the vision stimuli, we collected human ratings for our three image sets (240 images total) on a set of 11 stimulus features using an independent pool of n=206 online participants (n=17-20 per feature). Some features paralleled those used for sentences (e.g., composition, which can be thought of as parallel to grammaticality for sentences; and semantic coherence, which can be thought of as parallel to plausibility for sentences); others were commonly discussed features of images (e.g., expansiveness and the amount of visual clutter; (Berman et al., 2014; Conwell, Hamblin, et al., 2024; Iigaya et al., 2021; Intraub & Richardson, 1989; McMahon et al., 2023; Reece & Danforth, 2017; Rosenholtz et al., 2007; Tarhan & Konkle, 2020b) (**Figure 5C**, <u>Methods</u>).

Three features varied with the degree of representation agreement. Two features—*Visual Clutter* and *Image Aesthetics—*were specific to the visual stimuli, whereas *Valence* was measured for both sentences an images, but only varied with representation agreement for images (**Figure 5D**). The high-agreement images were rated as higher in visual clutter than the random images, which in turn, were rated higher than the low-agreement images (mean over participants: high-agreement: 0.10, random: ∼0, low-agreement:-0.11, *p*-value (high>random)=0.001, *p*-value (random>low)<0.001, paired-samples t-test). However, *Image Aesthetics* showed the opposite trend, with the high-agreement images rated as less aesthetically pleasing than the random images, which in turn, were rated lower than the low-agreement images (high-agreement:-0.10, random: ∼0, low-agreement: 0.10, *p*- value(high<random)=0.005, *p*-value(random<low)<0.001, paired-samples t-test). *Valence* showed a similar pattern to *Image Aesthetics* trend: high-agreement images were rated as more negative than the random images, which in turn, were rated lower than the low-agreement images (high-agreement:-0.07, random: 0.01, low-agreement: 0.06, *p*-value (high<random)=0.008, *p*-value (random<low)=0.043, paired-samples t-test). Finally, in contrast to sentences, we did not observe a reliable trend for *Perceived Frequency* in images: whereas the high-agreement images were rated as more frequent than the random images, this was not the case when comparing the random and the low-agreement images (high-agreement: 0.17, random:-0.08, low-agreement:-0.09, *p*-value(high>random)<0.001, *p*-value(random>low)=0.38, paired-samples t-test; see **Suppl. Figure 4B** for a related feature—*Image Typicality*—which showed a similar pattern to *Perceived Frequency*). Thus, in our initial observations, we identified some features that varied as a function of agreement for both sentences and images, but no features were shared between the language and vision domains.

## Discussion

In this study, we developed a method for computing the degree of representation agreement across artificial models (and humans) to test the *representation universality* hypothesis—the idea that the representational axes that are shared across high-performing artificial neural networks (ANNs) are the most aligned with the representational axes in human brains, plausibly determined by the input statistics of the relevant domain. For both the domains of language and high-level vision, we found that representations that are convergent across different models— and therefore more universal—align better with human neural representations, compared to representations that capture model-specific features (*representation individuality* hypothesis). In the domain of vision, we further showed that representation universality is bidirectional: stimuli whose representations are convergent across individual brains, as measured with fMRI, are also represented more similarly across ANNs. Finally, we found that the representational axes are largely distinct between language and vision: for language, the axes have to do with perceived sentence frequency and meaning generality, and for vision—with visual clutter, valence, and aesthetics.

Our results are consistent with recent work arguing for shared representational dimensions among vision ANNs, and between these ANNs and the human visual cortex (Z. Chen & Bonner, 2024). However, we show that representational convergence does not hold across the entire stimulus space (in either language or vision): instead, we find some parts of the space that show high convergence but other parts remain unaligned, i.e. idiosyncratic, across models and across individual brains. For ANNs, these idiosyncrasies may result from differences in architecture, objective function, learning algorithms, and other design choices; for humans, they may result from individual experiences with the relevant domain. The shared axes, on the other hand, likely reflect the structure of the relevant domain, which is robustly captured across model design choices and individual experiences.

Our findings are also in line with a recent proposal of representational convergence across ANNs (the *Platonic Representation* hypothesis): Huh et al. (2024) argue that the representations learned by high-performing ANNs are converging (within and across domains) because these models are modeling the same underlying world. We extend this general idea to biological networks. Importantly, however, although we find evidence of representation universality in two domains, language and vision, we make no claims about *across-domain* representational convergence. In fact, our behavioral rating study suggests that the coding axes differ between language and vision, which is perhaps to be expected given that our visual system is shaped by our perceptual experiences, whereas our language system necessarily abstracts away from the perceptual inputs focusing on the level of description that is suitable for communicating about the world.

The behavioral rating studies provide some initial hints about possible axes in the representations of sentences and images, and suggest that largely distinct features may organize the representational spaces for language and vision. Frequency was measured as a feature in both domains, but only showed a reliable effect from high-agreement to random to low-agreement sentences, not images. Instead, visual clutter, valence, and image aesthetics showed reliable effects for the visual stimuli. Nevertheless, future work may discover some features, perhaps simpler statistical features than the ones examined here, that would uncover similar organizing dimensions of the universal representational across language and vision (Goetschalckx et al., 2019; Graham & Field, 2008; Iigaya et al., 2021).

### Implication for computational neuroscience

A common approach in computational neuroscience has been to treat ANNs—distinguished by their architecture, task objective, or training data diet—as distinct hypotheses of how some biological system of interest (e.g., IT cortex or the language system) performs the relevant task (Doerig et al., 2023; Dong et al., 2018; Richards et al., 2019; Schrimpf et al., 2020; Yamins & DiCarlo, 2016; Yamins et al., 2014). Some large-scale efforts have even focused on systematically comparing large numbers of ANNs against a set of behavioral and neural benchmarks (Conwell, Prince, et al., 2024; Schrimpf et al., 2018; Willeke et al., 2022) with the goal of identifying the best model for the domain in question. Our results suggest a complementary approach of abstracting away from any particular ANN and focusing on representations that are similar (universal) *across* models (see also (Lindsay & Bau, 2023), given that such representations are most likely to reflect the structure of the input in relevant domain rather than model-specific features.

Our work also underscores the importance of careful stimulus selection in comparing models against the brain. Although the current tendency in the field to use naturalistic stimuli is justified on ecological validity grounds, it should not come at the expense of critically considering and evaluating alternative hypotheses, which may require sampling or designing stimuli that do not follow the natural distribution (Bowers et al., 2022; Gershman, 2021; Platt, 1964; Rust & Movshon, 2005). Only through careful selection of sentences and images from the aligned and misaligned parts of the representational space were we able to establish representation universality across ANNs and between ANNs and brains.

### Convergent evolution in biology and between artificial neural networks and brains

Conceptually, our work follows a comparative approach common in biology and neuroscience. Behaviors and neural mechanisms that are similar across species may be inherited from a common ancestor (homology), or may evolve independently, shaped by environmental and survival needs (analogy or convergence). Convergent traits across diverse species afford insights into why certain behaviors or neural mechanisms may be particularly useful and efficient. A notable example of a universal mechanism is the neural representation underlying spatial cognition (grid cells), initially discovered in rodents (Hafting *et al*., 2005), and subsequently identified in bats (Yartsev et al., 2011), monkeys (Killian et al., 2012), and humans (Jacobs *et al*., 2013). More recently, similar representations have been observed in task-optimized neural networks (Banino et al., 2018; Cueva & Wei, 2018). Whereas similarity across biological species (within a clade) might suggest a phylogenetically conserved mechanism, similarity between brains and ANNs clearly reflects environmentally-driven convergence: the need to solve a particular problem in the external world, be it navigation, or face recognition, or next word prediction (Cao & Yamins, 2024; Dobs et al., 2022; Kanwisher et al., 2023; Maheswaranathan et al., 2023; Moser, 2014; Turing, 1952; Turner et al., 2019; P. Y. Wang et al., 2021).

### Implications for AI

Uncovering universal principles shared across ANNs is a central focus in the mechanistic interpretability work in AI (Chughtai et al., 2023; Goh et al., 2021; Gurnee et al., 2024; Mehrer et al., 2020; Olah et al., 2020; Olsson et al., 2022). A common approach has been to use small neural network models to develop and test hypotheses about some aspect of the representations or computations (e.g., (Saxe et al., 2019), and then, in some cases, extend these tests to larger models to test for the generality of such representations (Chughtai et al., 2023; Elhage et al., 2022). Our work advocates a complementary approach based on distilling universal aspects of representation across ANNs. Working with such distilled representations is akin to working with small models in that many irrelevant (in this case, model-specific) features are removed, allowing for inferences that are challenging to make with large off-the-shelf models. Although previous studies have explored universal representations across ANNs (Bansal et al., 2021; Barannikov et al., 2021; Ding et al., 2021; Huh et al., 2024; Moor et al., 2020; J. Wang et al., 2018), our work further ties these representations to those found in biological neural networks: brains.

### Limitation and future work

Our results can be strengthened in several ways. First, we focused on representational similarity to find convergence across models due to its ease of interpretation and prior studies on alignment between ANNs and the brain (Khaligh-Razavi & Kriegeskorte, 2014; Yamins et al., 2014). However, we did not systematically evaluate which aspects of representations are detected by representational similarity. Our metric is based on the global geometry of representation, while recent work has used topological features, such as neighborhood relationship (Huh *et al*., 2024), or one-to-one comparison of axes of representation (Khosla *et al*., 2024) in comparing among ANNs, and between ANNs and neural representations. Second, constructing joint universal representations from the full richness of the stimulus space, instead of sub-sampling from the high-vs. low-agreement portions of the space, could provide a clearer picture of both convergent and divergent aspects of representations across ANNs, and result in greater convergence with human brains and behavior. Such a space may also facilitate the interpretation of the coding axes in both language and vision. Finally, the availability of a well-characterized shared representational space in any domain could serve as a tool for isolating components of the representations that are specific to particular individuals or groups of individuals, and may help understand how representations get warped by brain conditions or psychiatric illnesses.

## Acknowledgements

The authors thank Josh McDermott, Sam Gershman, Nancy Kanwisher, Niko Kriegeskorte, Tal Golan, Alex Williams, Andrew Lampinen, Chengxu Zhuang, Cory Shain, Martin Schrimpf and his group, Stanislas Dehaene and his group, and the audiences at CCN 2023 (at Oxford, U.K.) and Ascona 2024 (at Ascona, Switzerland) for helpful comments and discussions. EH, CC, NZ, MR, and EF were partially supported by NIH grant U01-NS121471.

CC was supported by a fellowship from the Kempner Institute for the Study of Natural and Artificial Intelligence at Harvard University. NZ was supported by a postdoctoral fellowship from the K. Lisa Yang Integrative Computational Neuroscience (ICoN) Center at MIT. EF was additionally supported by funds from MIT’s McGovern Institute for Brain Research, Quest for Intelligence, Department of Brain and Cognitive Sciences, and the Simons Center for the Social Brain.

## Methods

### 1. Stimulus selection

#### Language Stimuli

We selected a large and diverse set of sentences for sampling and optimization from the Universal Dependency (UD) corpus (**Table 1**(de Marneffe et al., 2021).The following steps were performed to filter out sentences:

1. Selection based on ASCII characters: only the following characters were allowed:

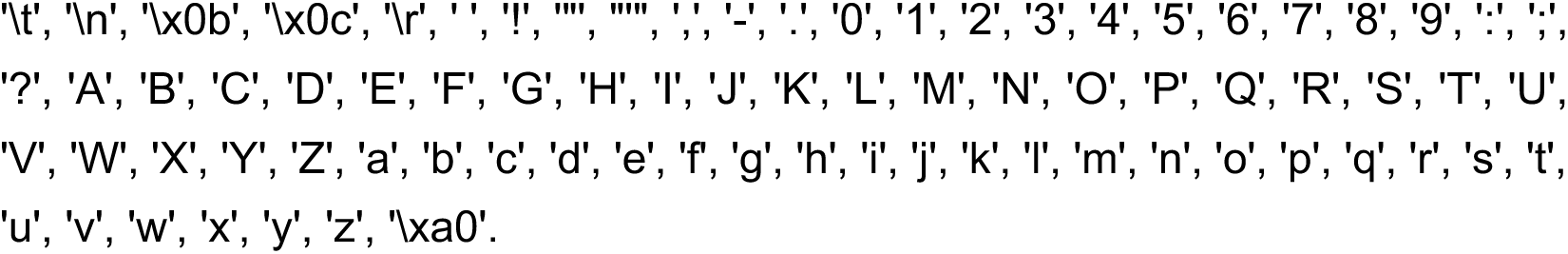
2. 2. Exclusion of sentences that contained: typos, abbreviations, foreign names, numbers (identified using the tags provided in the corpus).
3. Exclusion of sentences that are shorter than 5 words or longer than 20 words.
4. Exclusion of sentences with the single ‘’ character.
5. Exclusion of sentences that contain words with more than 3 uppercase letters.
6. Exclusion of duplicate sentences.

After filtering, 8409 sentences remained that were used for the sampling procedure designed to modulate representational agreement (see **Methods: stimulus optimization**).

In Experiment 1, we performed stimulus optimization over 8409 sentences. In Experiment 2, we wanted to further control the utterance length of the sentences, and did an additional filtering step based on the estimated length of the utterances. Specifically, we used text-to-speech models (Shen *et al*., 2017) to create an estimate of the utterance lengths for each sentence in the set of 8409, and filter out utterances with lengths less than 2 seconds or more than 4 seconds. The resulting set contained 5319 sentences.

**Table 1:** The genre composition of the Universal Dependencies English corpus (33,945 total sentences).

| Corpus | # Sentences | Genres |
| --- | --- | --- |
| UD_English-GUM | 5961 | Academic, fiction, news, nonfiction, spoken, web, wiki |
| UD_English-EWT | 16622 | Blog, email, reviews , social |
| UD_English-ESL | 5124 | Learner essays |
| UD_English-PUD | 1000 | News, wiki |
| UD_English-LinES | 5243 | Fiction, nonfiction, spoken |
| UD_English-GUMReddit | 895 | Blog, social |

#### Linguistic features

We extracted a set of lexical and syntactic features for each sentence to ensure that the sets of sentences (high-agreement, random, and low-agreement) were approximately matched across these features. For each word in a given sentence, we extracted (1) age of acquisition (Kuperman et al., 2012), (2) concreteness (Brysbaert et al., 2014), (3) prevalence (Brysbaert *et al*., 2019), (4) arousal (Warriner et al., 2013), (5) valence (Warriner et al., 2013), (6) ambiguity (i.e., number of meanings) (Brysbaert & New, 2009), (7) lexical frequency (Brysbaert & New, 2009), (8) 3-gram surprisal (Brants & Franz, 2009), and (9) word length (number of letters). For each sentence, we additionally extracted (10) length (number of words) (Tuckute *et al*., 2024). For features 1-9, we averaged the values across the words for which the measures were available to derive the value for the sentence. For feature 10 we obtained one value per sentence. We then used sentence values to compare linguistic features between full, random, and low-agreement sentence sets in Experiment 1 (**Figure 1E**, **Suppl. Figure 1B)**, and between high-agreement, random, and low-agreement sentence sets in Experiment 2 **(Suppl. Figure 2B)**.

#### Language models

We selected seven high-performing, transformer-based language models to use for selecting stimuli that were in high-agreement or low-agreement across models. These models were chosen to span a wide range of architectures, training datasets, and objective functions, and they were all shown to perform well in predicting fMRI responses in prior work (Schrimpf *et al*., 2021) (**Figure 1B**). For each model, we performed the stimulus sampling process using the model layer that best predicted fMRI responses on the Pereira 2018 benchmark (**Table 2**, see ‘Model-to-brain alignment’ below).

**Table 2:** The language models, and their corresponding layer, used in the stimulus sampling procedure.

| Language Model | Best Performing Layer |
| --- | --- |
| roberta-base | encoder.layer.1 |
| bert-large-uncased-whole-word-masking | encoder.layer.11.output |
| albert-xxlarge-v2 | encoder.albert_layer_groups.4 |
| xlm-mlm-en-2048 | encoder.layer_norm2.11 |
| xlnet-large-cased | encoder.layer.23 |
| Ctrl | h.46 |
| gpt2-xl | encoder.h.43 |

**BERT:** First introduced by (Devlin et al., 2018), BERT is a bidirectional transformer model trained on a combination of masked language modeling (MLM) and next sentence prediction objectives. This model utilized a training dataset comprising BookCorpus (Zhu et al., 2015) and English Wikipedia (Devlin *et al*., 2018). Inputs were tokenized using a byte-level Byte Pair Encoding (BPE) algorithm (Sennrich et al., 2015); see Devlin et al., 2018 for additional details about the model architecture and training). We used the ‘bert-large-uncased-whole-word-masking’ model configuration from the *Hugging Face* transformers library (Wolf et al., 2019), which has a hidden size of 1024, 24 layers, and 16 attention heads.

**RoBERTa:** First introduced by (Liu et al., 2019), the RoBERTa architecture is identical to that of BERT (see above). The primary training objective of RoBERTa is masked language modeling (MLM); however, the location of the mask was randomized ten times in the training dataset. The training dataset for the model includes a combination of BookCorpus (Zhu et al., 2015), CCNews (Nagel, 2016), OpenWebText (Gokaslan and Cohen, 2019), and STORIES (Trinh and Le, 2018). The input was tokenized using the WordPiece algorithm (see Liu et al., 2019 for additional details about the model architecture and training). We used the ‘roberta-base’ model configuration from the *Hugging Face* transformers library (Wolf et al., 2019), which has a hidden size of 768, 12 layers, and 12 attention heads.

**ALBERT:** First introduced by (Lan et al., 2019), the ALBERT architecture is based on the BERT architecture and incorporates several modifications to reduce the number of parameters in the model. Specifically, it employs factorized embedding, serving as an intermediate representation between the dictionary and the embedding layer, and it implements cross-layer parameter sharing. ALBERT is primarily trained with a MLM objective. However, unlike BERT, ALBERT uses a different secondary objective—self-supervised sentence order prediction instead of next sentence prediction. The training dataset is a combination of BOOKCORPUS (Zhu et al., 2015) and English Wikipedia (Devlin *et al*., 2018), similar to that used for BERT. Inputs were tokenized using a SentencePiece algorithm (Kudo & Richardson, 2018); see Lan et al., 2019 for additional details about the model architecture and training). We used the ‘albert-xxlarge-v2’ model configuration from the *Hugging Face* transformers library (Wolf et al., 2019), which has a hidden size of 4096, 12 layers, and 64 attention heads.

**XLM:** First introduced by (Lample & Conneau, 2019), XLM incorporates a cross-lingual objective combined with a MLM objective. The cross-lingual objective is similar to MLM, except it involves training with multiple languages in parallel. The training dataset is composed of English Wikipedia (Attardi, 2015). Inputs were tokenized using the BPE algorithm (Sennrich et al., 2015; see Lample & Conneau, 2019 for additional details about the model architecture and training). We used the ‘xlm-mlm-en-2048’ model configuration from the *Hugging Face* transformers library (Wolf et al., 2019), which has a hidden size of 2048, 12 layers, 16 attention heads, and is non-causal.

**XLNET:** First introduced by (Yang et al., 2019), XLNet is an autoregressive transformer model with the objective of maximizing the expected likelihood of a sequence given all permutations of the input context (for n items: n! permutations). Although it features an autoregressive transformer architecture, it incorporates a recurrence mechanism from earlier models (Dai et al., 2019). The original model was trained on multiple datasets including BookCorpus (Zhu et al., 2015), English Wikipedia (Devlin *et al*., 2018), Giga5 (Parker et al., 2011), ClueWeb2012-B (Callan *et al*., 2009), and Common Crawl (Nagel, 2016). Inputs were tokenized using the SentencePiece algorithm (Kudo & Richardson, 2018); see Yang et al., 2019 for additional details about the model architecture and training). We used the ‘xlnet-large-cased’ model configuration from the *Hugging Face* transformers library (Wolf et al., 2019), which has a hidden size of 768, 24 layers, and 16 attention heads.

**CTRL:** First introduced by (Keskar et al., 2019), CTRL is an autoregressive transformer model that incorporates an additional code variable appended to the context. It was trained using a causal language modeling (CLM) objective. The training dataset is composed of a diverse collection of texts including Wikipedia, Project Gutenberg (Project Gutenberg, n.d.), Reddit, OpenWebText (Gokaslan and Cohen, 2019), news dataset (Hermann *et al*., 2015), Amazon reviews (McAuley *et al*., 2015), WMT (Barrault *et al*., 2019), ELI5 (Fan *et al*., 2019), MRQA (Fisch *et al*., 2019), and a few other datasets. Inputs were tokenized using the BPE algorithm (Sennrich et al., 2015); see Keskar et al., 2019 for additional details about the model architecture and training). We used the ‘ctrl’ model configuration from Hugging-Face transformers library (Wolf et al., 2019), which has a hidden size of 1280, 48 layers, and 15 attention heads.

**GPT-2:** First introduced by (Radford et al., 2019), GPT-2 is an autoregressive transformer model trained on a CLM objective. It was trained using the WebText dataset (Radford *et al*., 2019), which notably excludes any Wikipedia pages. The inputs were tokenized using a BPE algorithm (Sennrich et al., 2015) (see Radford et al., 2019 for additional details about the model architecture and training). We used the ‘gpt2-xl’ model configuration from the *Hugging Face* transformers library (Wolf et al., 2019), which has a hidden size of 1280, 48 layers, and 15 attention heads.

The following models were not used in the stimulus sampling procedure but instead served (i) to test the generalizability of our findings to smaller models in the same model class (DistilGPT2, also ‘gpt2-small’, ‘gpt2-medium’, and ‘gpt2-large’ described below, **Figure 2D, Suppl. Figure 3**), (ii) to test the generalizability of our findings to newer, larger models (LLAMA-7B and LLAMA-68B, **Figure 2E**), and (iii) as a control to test whether the critical effects were present in non-transformer-based models (Word2Vec and LM-1B, Supplementary Figure 3) or a random embedding space (Supplementary Figure 3).

**DistilGPT2:** First introduced by (Sanh *et al*., 2019), distilGPT2 is a smaller version of GPT-2 architecture (82 million parameters—around 70% of the number of parameters as GPT-2 small).

The model was trained on a distillation objective, where the smaller model (DistilGPT2) learns to mimic the teacher model (GPT-2). Inputs were tokenized using the BPE algorithm (Sennrich et al., 2015); see Sanh et al., 2019 for additional details about the model architecture and training). We utilized the ‘distilgpt2’ configuration from the *Hugging Face* transformers library (Wolf *et al*., 2019) which has a hidden size of 768, 6 layers, and 12 attention heads.

**GPT2 additional models:** We used three additional models with the same architecture and training as ‘gpt2-xl’, but with varying model sizes (Radford *et al*., 2019).

- ‘gpt2-small’: Hidden size = 768, 12 layers, 12 attention heads; 117 million parameters.
- ‘gpt2-medium’: Hidden size = 1024, 24 layers, 16 attention heads; 345 million parameters.
- ‘gpt2-large’: Hidden size = 1280, 36 layers, 20 attention heads; 774 million parameters.

**LLAMA:** First introduced by (Touvron *et al*., 2023), LLAMA is an autoregressive transformer model trained on a CLM objective. It was trained using a diverse compilation of sources, including English Common Crawl (Touvron et al., 2023), C4 (Raffel *et al*., 2019), GitHub, English Wikipedia (Devlin *et al*., 2018), Project Gutenberg (Project Gutenberg, n.d.), Book3 (Touvron *et al*., 2023), ArXiv (Touvron et al., 2023), and StackExchange (Touvron *et al*., 2023). Inputs were tokenized using the BPE algorithm (Sennrich et al., 2015); see Touvron et al., 2023 for additional details about the model architecture and training). We utilized two variants of the architecture selected from the *Hugging Face* library (Wolf et al., 2019):

- ‘LLAMA-7B’: Hidden size = 4096, 32 layers, 32 attention heads; 6.7 billion parameters.
- ‘LLAMA-68B’: Hidden size = 8192, 80 layers, 64 attention heads; 65.2 billion parameters.

**Word2Vec:** First introduced by (Mikolov *et al*., 2013; Mikolov *et al*., 2013), Word2Vec is an embedding model trained to predict words based on the surrounding context. It was trained on a Google News dataset containing 100 billion words (see Mikolov et al., 2013; Mikolov et al., 2013 for additional details about the model and training). We utilized a word-level tokenizer (Wolf *et al*., 2020) and defined the sentence embedding as the average embedding over all words in the sentence.

**LM-1B:** First introduced by (Jozefowicz *et al*., 2016), LM-1B is a Long Short-Term Memory (LSTM) model with two layers that was trained to perform next word prediction on 1 billion words (Chelba et al., 2014; see Jozefowicz et al., 2016 for additional details about the model and training). Inputs were tokenized using the BPE algorithm (Sennrich et al., 2015).

**Random Embedding:** We used a random embedding as a control model with an embedding dimension of 1600, similar to GPT2-xl. For each sentence, we drew an independent sample from a uniform distribution with a span of [0,1).

#### Vision Stimuli

We used the subset of stimuli from Microsoft Common Object in Context (COCO) dataset (Lin *et al*., 2014) that were overlapping across participants in the Natural Scenes Dataset (NSD) fMRI dataset (Allen *et al*., 2022). COCO dataset contains photographs gathered from online repositories, but NSD dataset stimuli were restricted to 2017 train/val splits (Lin *et al*., 2014), with pixel-level annotation of ‘stuff’, such as sky and land, and ‘things’ such as car and hat (see (Allen et al., 2022) for more details).

#### Vision Models

We selected seven high-performing, transformer-and convolution-based vision models to use for selecting stimuli that were in high-agreement or low agreement across models. These models were chosen to span a wide range of architectures, training datasets, and objective functions, and they were all shown to perform well in predicting fMRI responses in prior work (Conwell et al., 2024). For each model, we performed the stimulus sampling process using the model layer that best predicted fMRI responses on the NSD occipito-temporal cortical parcels (OTC) to be consistent with (Conwell, Prince, et al., 2024) (**Table 3**, see ‘Model-to-brain alignment’ below).

**Table 3:**
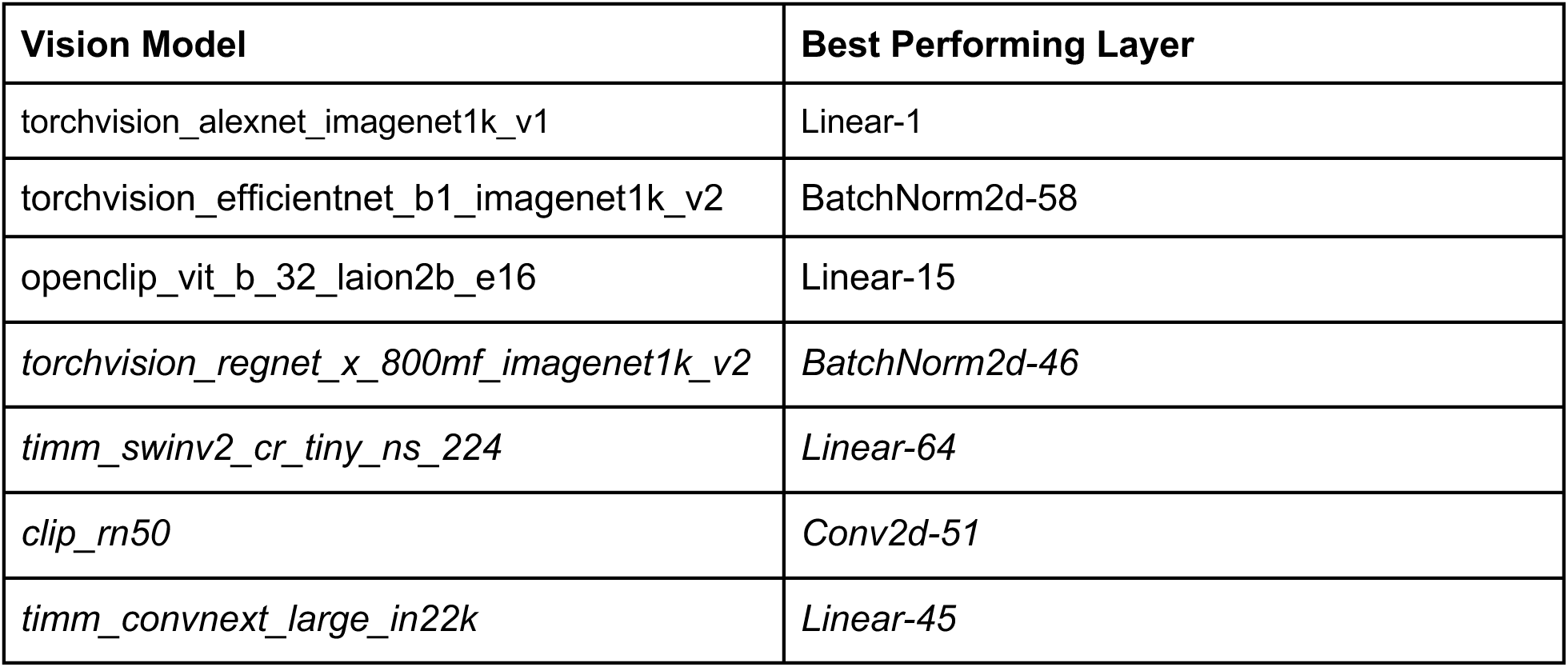
The vision models, and their corresponding layer used in the stimulus sampling procedure.

**Table 4:**
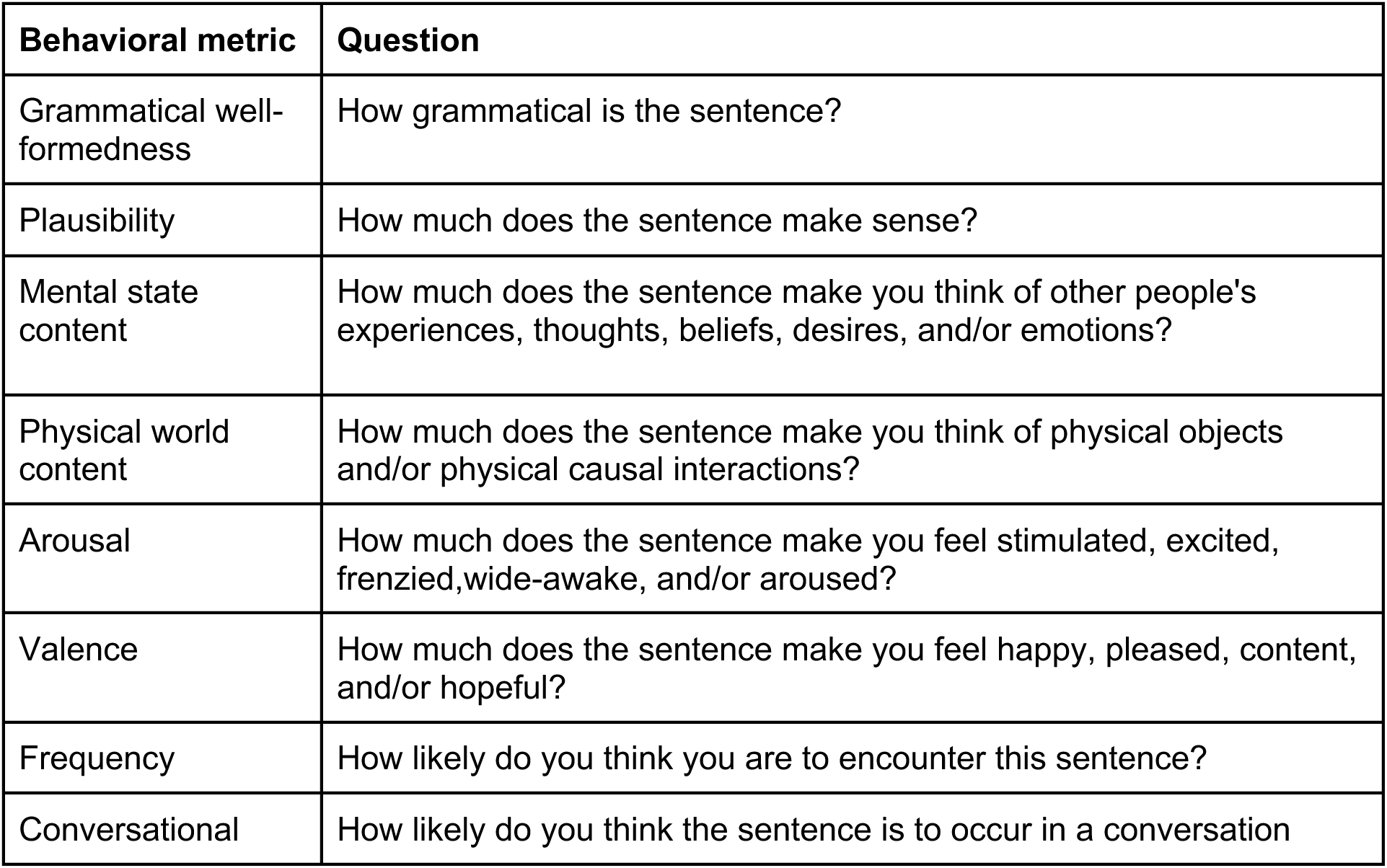
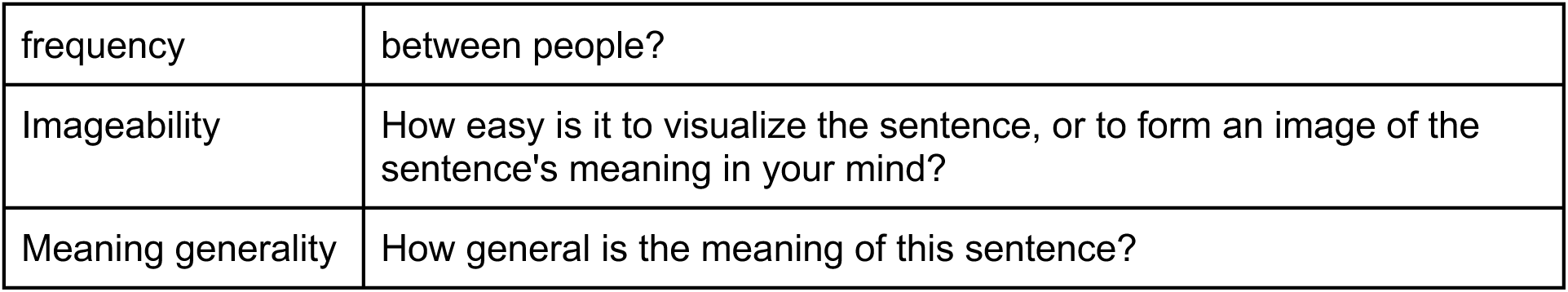
Behavioral questionnaires for stimuli used in Experiment 2.

**AlexNet:** First introduced by (Krizhevsky et al., 2017), AlexNet is a convolutional neural network (CNN) trained on the ImageNet1K_v1 dataset (Russakovsky *et al*., 2015) using a supervised classification objective function (see Krizhevsky et al., 2017 for additional details about the model architecture and training). We used the ‘torchvision_alexnet_imagenet1k_v1’ model from the *PyTorch torchvision* library.

**EfficientNet**: First introduced by (Tan and Le, 2019), EfficientNet is a CNN trained on the ImageNet1K_v2 dataset (Russakovsky *et al*., 2015) using a supervised classification objective function (see Tan and Le, 2019 for additional details about the model architecture and training). We used the ‘torchvision_efficientnet_b1_imagenet1k_v2’ model from the *PyTorch torchvision* library.

**ViT**: First introduced by (Radford et al., 2021a), ViT is a multimodal vision-language transformer neural network trained on the LAION2B dataset (Schuhmann *et al*., 2022); see Radford et al., 2021a for additional details about the model architecture and training). We used the ‘openclip_vit_b_32_laion2b_e16’ model from the *OpenCLIP* library (Ilharco et al., 2021).

**RegNet**, First introduced by (Radosavovic et al., 2020), RegNet is a CNN trained on the ImageNet1K_v2 dataset (Russakovsky *et al*., 2015) using a supervised classification objective function (see Radosavovic et al., 2020 for additional details about the model architecture and training). We used the ‘torchvision_regnet_x_800mf_imagenet1k_v2’ model from the *PyTorch torchvision* library.

**Swin:** First introduced by (Liu et al., 2021), Swin is a transformer neural network trained on the ImageNet1K dataset (Russakovsky *et al*., 2015) using a self-supervised objective (see Liu et al., 2021 for additional details about the model architecture and training). We used an independent implementation of the model ‘timm_swinv2_cr_tiny_ns_224’ model from the *PyTorch timm vision* library.

**CLIP**: First introduced by (Radford et al., 2021b), CLIP is a hybrid convolutional vision-language model by OpenAI trained on 400 million internet-based image/text pairs using a contrastive objective (see Radford et al., 2021b for additional details about the model architecture and training). We used the ‘clip_rn50’ model from the *OpenCLIP* library (Ilharco et al., 2021).

**ConvNext**: First introduced by (Liu et al., 2022), ConvNext is a CNN trained on both the ImageNet1k and ImageNet21k datasets (Russakovsky *et al*., 2015) using a supervised classification objective (see Liu et al., 2022 for additional details about the model architecture and training). We used the ‘timm_convnext_large_in22k’ model from the *PyTorch timm* library.

#### Stimulus optimization

**Language stimuli.** To sample a set of sentences for which models disagree (the low-agreement set), we used a coordinate ascent algorithm (Boyd & Vandenberghe, 2004) to search over the full set of curated N sentences extracted from the Universal Dependency corpus to find a set sentences with size *n* that maximized/minimized the correlation distance between the RDMs of pairs of models.

For each model, we fed the full set of *N* sentences as the input from the first word to the last word after tokenization specific to each model (excluding the period). The sentence-level representation was defined as the activations at the last word (function *readwords* in the *neural-nlp* pipeline, adopted from Schrimpf et al., 2021). Given a sample set of sentences *S_n_*, at each step of the algorithm, we computed the representational dissimilarity matrix (RDM(Nili et al., 2014) between sentence-level representations in *S_n_* using Pearson correlation coefficient (*M*_i_[*S*_n_] for model *i*). We then defined model pair disagreement between model *M_i_* and model *M_j_* as:

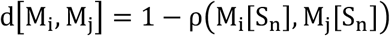

where ρ is Pearson’s correlation coefficient. The model agreement *a_M_* for all model pairs *M_i_* a *M_j_* was computed as:

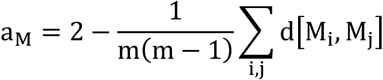

where *m* is the number of models. The set of sentences *S_n_* was selected through iterative substitution from the full set of sentences *S_N_*. In each step of the optimization loop, individual sentences were swapped with new ones from *S_n_* and a new *a_M_* was computed; if the value of *a_M_* improved in the desired direction by substitution (increase for Experiment 1; increase or decrease for Experiment 2), then the swap was considered successful. This process was performed over all sentences in *S_n_* in each iteration of the algorithm and was repeated until there was no swapping effective in changing *a_M_*. We note that depending on the initial choice of *S_n_*, the optimization algorithm can, in principle, arrive at multiple solutions that maximize/minimize agreement to the same extent.

**Language-Experiment 1:** to find low-agreement sentence set, we implemented an early stopping algorithm given the computational cost of iteration over all sentences. We performed 5 optimizations with different initial *S_n_* each resulting in a set of 300 optimized sentences. We then manually filtered out sentences that contained offensive and sensitive content. To get to the final 200 sentences, we started by selecting sentences that were shared across all 5 stimulus sets, then selecting sentences that were shared across 4 stimulus sets, and so on, until we reached 200 sentences. We then recomputed *a_M_* for this new set of sentences to ensure that representational agreement was close to the agreement on the original optimized sets.

**Language-Experiment 2**: here we removed the early stopping criterion and instead iterated over the full sentence set to find the high-agreement and low-agreement sentence sets. We achieved this by implementing the algorithm on a GPU and precomputing all sentence dissimilarities across models. We additionally reduced the size of the sentence set we searched over from 8409 to 5319, to have more control over utterance length. The optimization resulted in a set of 100 sentences that either maximized or minimized representational agreement across models. To maximize agreement we used the same *a_M_* as in Experiment 1. To minimize agreement instead we used 2 – *a_M_* as our measure. We then manually filtered out sentences that contained offensive or sensitive content, resulting in 80 final sentences in each set. We subsequently recomputed *a_M_*and 2 – *a_M_*for these new sets to ensure the agreement values were similar to those of the original optimized sets.

To derive a random set of sentences, we extracted 1000 sets of 100 randomly drawn sentences from the full set of N=5319 sentences. We then selected the set that was closest to the mean representational agreement across all 1000 random sets. Finally, we manually filtered out sentences that contained offensive or sensitive content, resulting in 80 final sentences.

**Vision-Experiment 3.** For the vision dataset (Allen et al., 2022), we followed the same optimization procedure as language, except an additional term was introduced to ensure that the *distribution* of stimulus dissimilarities was not significantly altered during the sampling process. We first computed an average distribution of image dissimilarity for 80 samples for each model (by sampling 1000 random sets of 80 images and computing an average distribution) and denoted each resulting average distribution as P_Mi_. For any random set of images, we measured the distance between sentence dissimilarities and P_Mi_ using KL divergence, denoted as 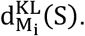. The range of 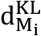 values could vary between models, but we wanted a single penalty to add to the original definition of agreement *a_M_*, so that the sampling procedure remains the same. To resolve this, we defined a normalization term as 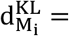 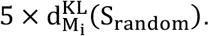 At each step of sampling, given the current sentence set *S_n_* we computed 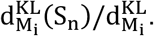 We then summed the normalized KL divergence values across all models and calculated the average d^KL^ If the d^KL^ was larger than threshold = 0.06, we then added it as a penalty to agreement *a_M_*. We experimented with several threshold values, to arrive at a value that achieves both changes in agreement measure and minimally affects the distribution of image dissimilarities between each set. Finally, we used a same approach as **Language-Experiment 2** to find random set of images.

**Vision-experiment 4:** here we modified the representation on which we performed sampling from ANNs to participants fMRI responses. Specifically we computed RDM of fMRI responses in OTC in each participants over the 1000 images (Allen et al., 2022). We then search over participants RDMs to find 3 sets of images with high-, random, and low-agreement. We used the same sampling approach as **Language-Experiment 2**, except that the agreement was computed and optimized over 6 participant-pairs (4 participants). Finally, we used a same approach as **Language-Experiment 2** to find random set of images.

## 2. fMRI data collection, preprocessing, and analysis

### Language Experiments

#### Language localizer

The regions of the language network were localized using a task described in detail in (Fedorenko *et al*., 2010) and subsequent studies from the Fedorenko lab (the task is available for download from https://evlab.mit.edu/funcloc/). Briefly, participants silently read sentences and lists of unconnected, pronounceable nonwords (each 12 word-/nonwords-long) in a blocked design. The sentences > nonwords contrast targets brain regions that that support high-level language comprehension. This contrast generalizes across tasks (Fedorenko et al., 2010; Ivanova et al., 2020; Scott et al., 2017) and presentation modalities (reading vs. listening; e.g. (X. Chen et al., 2023; Fedorenko et al., 2010; Malik-Moraleda et al., 2022; Scott et al., 2017). All the regions identified by this contrast show sensitivity to lexico-semantic processing (e.g., stronger responses to real words than nonwords) and combinatorial semantic and syntactic processing (e.g. stronger responses to sentences and Jabberwocky sentences than to unstructured word lists and nonword lists) (Blank et al., 2016; Fedorenko et al., 2020, 2010, 2012, 2016; Shain et al., 2024). More recent work further shows that these regions are also sensitive to sub-lexical regularities (Regev *et al*., 2024), in line with the idea that this system stores our linguistic knowledge, which encompasses regularities across representational grains, from phonological and morphological schemas to words and constructions (see (Fedorenko et al., 2024) for a review). Stimuli were presented one word/nonword at a time at the rate of 450 ms per word/nonword. Participants read the materials passively and performed a simple button-press task at the end of each trial, which was included in order to help participants remain alert. Each participant completed 2 ∼6 min runs.

#### Experiment 1 design and procedure

Participants listened to the auditory recordings of the 200 sentences selected as described in **Methods: Stimulus Optimization**, along with 16 sets of nonword sequences, which were included as a baseline. The sentences and nonwords were recorded by a native English speaker and the length was adjusted using custom-written code in MATLAB software (by stretching or compressing the audio while preserving the pitch, plus optional additional padding) so that the duration of each stimulus was a multiple of the TR value (2 s). The final distribution of sentence lengths was as follows: 26 sentences - 2 s, 114 sentences - 4 s; 75 sentences - 6 s, 2 sentences - 8 s; all nonword sequences were 4 s. The stimuli were divided into four sets, which corresponded to scanning runs. Each set consisted of 50 sentences and 8 nonword sequences each, plus, two of the sentences were repeated within the run to allow for the measurement of response reliability within the run. The stimuli were presented with an inter-trial interval of 4 s, and each run additionally included three longer fixation periods of 12 s. The runs lasted 8.4 minutes on average, varying between 8 and 9 minutes. Participants completed 8 runs (2 repetitions of each set) on the first day of the experiment, and a second set of 8 runs on a different day (5-13 days after the first visit). Thus, across the two scanning sessions, each sentence was presented 4 times. However, because for language stimuli repetitions beyond 2 instances do not appear to improve, and often decrease, response reliability (unpublished data from the Fedorenko lab), we used the data from the first session only, except for one participant, for whom the data in the second session were higher quality.

#### Experiment 2 design and procedure

Participants listened to the auditory recordings of the 240 sentences selected as described in **Methods: Stimulus Optimization**. The sentences were recorded by a native English speaker, and the length was adjusted using custom-written code in MATLAB software (by stretching or compressing the audio while preserving the pitch, plus optional additional padding) so that the duration of each stimulus was 2 TRs (4 s). The stimuli were divided into four sets, which corresponded to scanning runs. Each set consisted of 60 sentences, and contained 20 sentences from high-agreement, random, and low-agreement groups in random order. The stimuli were presented with an inter-trial interval of 4 s, and each run additionally included three longer fixation periods of 12 s. The runs lasted 8.4 minutes. The participants listened to 2 repetition of each set over the course of ∼2 hour scanning session, in palindromic order (example: sets: [1,2,3,4,4,3,2,1]) and we further shuffled the order of sentences between first and second repetitions of each set.

#### fMRI data acquisition

Structural and functional data were collected on a whole-body 3 Tesla Siemens Prisma scanner with a 32-channel head coil at the Athinoula A. Martinos Imaging Center at the McGovern Institute for Brain Research at MIT. T1-weighted, Magnetization Prepared Rapid Gradient Echo (MP-RAGE) structural images were collected in 208 sagittal slices with 0.85 mm isotropic voxels (TR = 1,800 ms, TE = 2.37 ms, TI = 900 ms, flip = 8 degrees). Functional, blood oxygenation level-dependent (BOLD) data were acquired using an SMS EPI sequence with a 90° flip angle and using a slice acceleration factor of 2, with the following acquisition parameters: fifty-two 2 mm thick near-axial slices acquired in the interleaved order (with 10% distance factor), 2 mm × 2 mm in-plane resolution, FoV in the phase encoding (A >> P) direction 208 mm and matrix size 104 × 104, TR = 2,000 ms, TE = 30 ms, and partial Fourier of 7/8. The first 10 s of each run were excluded to allow for steady state magnetization

#### fMRI preprocessing

fMRI data were analyzed using SPM12 (release 7487), CONN EvLab module (release 19b), and other custom MATLAB scripts. Each participant’s functional and structural data were converted from DICOM to NIFTI format. All functional scans were coregistered and resampled using B-spline interpolation to the first scan of the first session (Friston *et al*., 1995). Potential outlier scans were identified from the resulting subject-motion estimates as well as from BOLD signal indicators using default thresholds in CONN preprocessing pipeline (5 standard deviations above the mean in global BOLD signal change, or framewise displacement values above 0.9 mm; (Nieto-Castanon, 2020). Functional and structural data were independently normalized into a common space (the Montreal Neurological Institute [MNI] template; IXI549Space) using SPM12 unified segmentation and normalization procedure (Ashburner & Friston, 2005) with a reference functional image computed as the mean functional data after realignment across all timepoints omitting outlier scans. The output data were resampled to a common bounding box between MNI-space coordinates (-90,-126,-72) and (90, 90, 108), using 2mm isotropic voxels and 4th order spline interpolation for the functional data, and 1mm isotropic voxels and trilinear interpolation for the structural data. Last, the functional data were smoothed spatially using spatial convolution with a 4 mm FWHM Gaussian kernel.

#### fMRI first-Level Analysis

Responses in individual voxels were estimated using a General Linear Model (GLM) in which each experimental condition (for the localizer data) or each stimulus (for the critical experiments) was modeled with a boxcar function convolved with the canonical hemodynamic response function (HRF) (fixation was modeled implicitly, such that all timepoints that did not correspond to one of the conditions/stimuli were assumed to correspond to a fixation period). Temporal autocorrelations in the BOLD signal timeseries were accounted for by a combination of high-pass filtering with a 128 seconds cutoff, and whitening using an AR(0.2) model (first-order autoregressive model linearized around the coefficient a=0.2) to approximate the observed covariance of the functional data in the context of Restricted Maximum Likelihood estimation (ReML). In addition to experimental condition effects, the GLM design included first-order temporal derivatives for each condition/stimulus (included to model variability in the HRF delays), as well as nuisance regressors to control for the effect of slow linear drifts, subject-specific motion parameters (6 parameters), and potential outlier scans (identified during preprocessing as described above) on the BOLD signal.

#### Identification of language-responsive regions

We defined a set of functional regions of interest (fROIs) using group-constrained, participant-specific localization (Fedorenko et al. 2010). For the core language fROIs, each individual map for the “sentences > nonwords” contrast was intersected with a set of 12 binary masks (6 in left hemisphere, 6 in right hemisphere). These masks (available for download from https://evlab.mit.edu/funcloc/) were derived from a probabilistic activation overlap map for the same contrast in a large independent set of participants (n = 220) using watershed parcellation, as described in (Fedorenko *et al*., 2010) for a smaller set of participants. These masks included 3 in the left frontal cortex—in the inferior and middle frontal gyri—and 3 in the left temporal and temporo-parietal cortex. Within each mask, a participant-specific language fROI was defined as the top 10% of voxels with the highest t-values for the localizer contrast (see (Lipkin et al., 2022) for evidence that the fROIs defined in this way are similar to the fROIs defined based on a fixed statistical significance threshold).

We evaluated the reliability of voxel responses in language regions to sentence stimuli. In Experiment 1, for each voxel we computed a correlation between responses to the first and second repetitions of the 200 sentences. We then compared these correlation values against a null distribution, which was created by shuffling the order of sentences and recomputing the correlation 500 times. A voxel was considered to show a significantly reliable response if the true correlation was higher than 95% of the shuffled correlations (i.e., 475 or more of the 500 correlations). Using this criterion, 3638 of the language-responsive voxels (33.4%) showed significant reliability.

In Experiment 2, the procedure was similar, except that the correlations were computed over the 80 sentences in the random set (to avoid potential biases toward voxels with differing reliability for the high-agreement or low-agreement sets). Using this approach 1702 of the language-responsive voxels (10.4%) showed significant reliability. The lower percentage of reliability prompted us to perform model-to-brain alignment on voxel responses to individual repetitions rather than averaging voxel responses across two repetitions (see below, Experiment 2-encoding model).

Finally, for each experiment, we selected voxels for each participant that were in the top 10% of repetition correlation values, while excluding voxels that did not pass the significance criterion described above. In Experiment 1, this selection process yielded [137, 135, 137, 137, 137, 137, 136, 137], totaling 1093 voxels across 8 participants. In Experiment 2, this process yielded [72, 136, 73, 43, 136, 136, 37, 127, 79, 100, 39, 108], totaling 1,086 voxels across 12 participants.

#### Vision experiment

We used the publicly available dataset of responses in occipitotemporal cortex (OTC) in humans to images of natural scenes, collected using 7-Tesla fMRI (Allen et al., 2022).

#### Experiment 3 design and procedure

Participants viewed images from the NSD dataset over 30–40 sessions using 7 Tesla fMRI acquisition, with approximately one scan session per week per participant. We specifically used data from four participants (sub01, sub02, sub05, and sub07, referred to as the group ‘Shared1000’ in (Allen *et al*., 2022). The images were displayed using a mirror system that projected images from an LCD monitor into the participants’ field of view (measured at 8.4° x 8.4°). Each image was presented for 3 s, followed by a 1s inter-trial interval period. There were 3 repetitions of images over the course of the experiment. Participants were asked to indicate whether the displayed image in a trial was new by pressing a button with the index finger of their right hand. Each run consisted of 63 trials, with 12s fixation period at the beginning, 16s at the end, and 5 fixation trials interleaved with stimulus trials. Each run lasted 5 minutes, and each session contained 12 runs.

#### fMRI data acquisition, preprocessing, and first-level analysis

We used the average voxel responses over the repetitions of each image in the ‘Shared1000’ portion of the NSD dataset for our analysis. The magnitude of responses for each voxel was calculated using the GLMSingle toolbox (Prince et al., 2022). Voxels were selected based on “noise ceiling signal-to-noise” values (NCSNR > 0.2). Details of fROI voxel selection are provided elsewhere (Conwell, Prince, et al., 2024). Across 4 participants, we analyzed 29480 voxels, [sub1:8088, sub2:7528, sub5:8015, sub7: 5849].

## 3. Model-to-brain comparisons

### Ceiling computation

Due to intrinsic noise in biological measurements, we estimated a ceiling value to reflect how well the best possible model of an average human could perform. Following (Schrimpf *et al*., 2021), we first subsampled—for each dataset separately— the data with n recorded participants into all possible combinations of s participants for all ∈ [2, n] (e.g. {2, 3, 4, 5, 6, 7,8} for Experiment 1). For each subsample s, we then designated a random participant as the target that we attempt to predict from the remaining *s* − 1 participants (e.g., predict 1 subject from 1 (other) subject, 1 from 2 subjects, 1 from 4,…, to obtain a mean score for each voxel in that subsample. For computational efficiency, we limit the number of combinations to 100 when choosing participants from a pool of n participant (for 12 participants there are 792 ways to draw 5 participants). To extrapolate to infinitely many humans and thus to obtain the highest possible (most conservative) estimate, we fit the equation 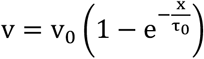 where *x* is each subsample’s number of participants, *v* is each subsample’s correlation score and *v_0_* and τ*_0_* are the fitted parameters for asymptote and slope respectively. This fitting was performed for each voxel independently with 200 bootstraps each to estimate the variance where each bootstrap draws *x* and *v* with replacement. The final ceiling value was the median of the per-voxel ceilings *v_0_*.

In Experiment 1, we first computed the average voxel response over 2 repetitions of sentences, and then computed the ceiling value per voxel. We then computed the median over voxels in each participant to get a participant level ceiling and computed the population ceiling by taking the median value over participants ceilings, and the final ceiling was estimated at 0.436 across the 8 participants. In Experiment 2, we estimated the ceiling over voxel responses over each repetition of stimuli separately, given that we did the model-to-brain mapping over individual repetitions. The estimated ceiling values for the first and second repetition were [0.251, 0.238] for high-agreement, [0.215, 0.197] for random, and [0.218, 0.281] for low-agreement.

#### Experiment 1, encoding model

To make direct comparison with results from (Schrimpf et al., 2021) we followed the same recipe and tested how well the model representation could predict human fMRI responses. To generate predictions, we used 80% of the sentences to fit a linear regression from the corresponding 80% of model representations to the corresponding 80% of fMRI responses. We applied the regression on model representations of the held-out 20% of stimuli to generate model predictions, which we then compared against the held-out 20% of fMRI responses with a Pearson correlation. This process was repeated five times, leaving out different 20% of stimuli each time, and we computed the per-voxel mean predictivity across those five splits. We aggregated these per-voxel scores by taking the median of scores for each participant’s voxels and then computing the median and median absolute deviation (m.a.d.) across participants (over per-participant scores). A reliable positive Pearson correlation value indicates that the model representation is able to predict fMRI responses with a linear transformation.

#### Experiment 1, RSA

Following (Schütt *et al*., 2023), we first computed a set of representational dissimilarity matrices (RDMs) for participants based on voxel responses. For each participant, we computed two RDMs—one per repetition—and averaged them for the final comparison with RDMs from models. Similarly, for each model, we computed an RDM based on layer activations. For both participants and models, RDMs were computed using correlation distance. We then compared model RDMs to participant RDMs using Pearson correlation. For two vectorized RDMs, r*_1_* and *r_2_*, Pearson correlation is defined as:

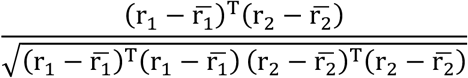

We also performed a model-to-brain RDM comparison using Spearman’s rho correlation (Schütt *et al*., 2023) and did not observe any qualitative differences in the results.

Similar to the encoding modeling, we computed a ceiling for the RSA metric using leave-one-out cross-validation (**Figure 2C**). Briefly, we built an RDM from *n* − 1 participants and compared it to the RDM of the remaining participant using the same Pearson correlation formula as above. This process was repeated for each participant, leaving one participant out at a time, and we reported the mean ceiling value across all eight participants.

#### Experiment 2, encoding model

Here, we employed a similar least-squares regression as in Experiment 1, albeit with some modifications. We observed that in Experiment 2, voxel reliability across participants was lower compared to the Experiment 1 dataset (33.4% of voxels across participants showed significant reliability over repetitions in Experiment 1, whereas in Experiment 2, this value was 10.4%). To mitigate the effect of low voxel reliability, we did not perform regression on the average voxel response. Instead, for each voxel, we computed an independent regression on each individual repetition of sentence sets (high-agreement, random, low-agreement). We then averaged the Pearson correlation values for each voxel over the two repetitions to arrive at a voxel-level prediction. Performing regression on individual repetitions allowed us to double the number of parameters used in model-to-brain alignment. Next, we took the median of voxel predictions to obtain participant-level predictions and subsequently took the median of participant predictions to derive population-level predictions, similar to the encoding model in Experiment 1. For each model, we constructed three regressions—one for each sentence group (high, random, low). The details of the regression are identical to those in the encoding model used in Experiment 1.

#### Experiment 3, encoding model

We followed a similar approach to that described in (Conwell et al., 2023) and performed a two-step regularized regression between ANN models and voxel responses across the OTC. First, we performed dimensionality reduction via sparse random projection to normalize the number of dimensions across models, as some vision models had a very large number of dimensions per image. The sparse random projection implementation we used was based on Scikit-learn (Pedregosa et al., 2011), which utilizes the Johnson-Lindenstrauss (JL) lemma (Achlioptas, 2001) to project n samples into a lower dimension while preserving their Euclidean distance up to a factor of 1 ± ɛ. We chose ɛ = 0.1, which resulted in 5920 dimensions.

To compute the alignment between representations in ANN vision models and voxels in the OTC, we implemented cross-validated ridge regression using the SciKit-learn RidgeCV tool, with alpha parameters ranging from 1 × 10^-1^ to 1 × 10^-7^. The best alpha for each voxel was selected based on the Pearson correlation between the predicted and actual responses (Conwell, Prince, et al., 2024). Using the optimal alpha, we performed 5-fold cross-validated regression for each voxel, splitting the data into 80% training and 20% testing for each fold. We then computed the average Pearson correlation between predicted and actual voxel responses across all folds. For each subject, we calculated the median voxel predictivity for each region and reported the median and median absolute deviation across the four subjects as the final metric, similar to the approach used in encoding modeling for comparing ANN language models with voxel activation in language regions.

#### Experiment 4, encoding model

We followed the same approach as **Experiment 3, encoding model** to calculate how well ANN vision models predicted voxel responses to image sets that were optimized over participants.

#### Linear mixed-effect modeling

To determine whether the differences observed in the encoding modeling results from Experiment 2 and Experiment 3 could be attributed to inter-stimulus similarity within individual models, we employed linear mixed-effects (LME) models (Seabold & Perktold, 2010).

Specifically, we used nested LME models to predict the Pearson correlation values obtained from the encoding analyses with either inter-stimulus similarity (a continuous predictor) or both inter-stimulus similarity and set (a treatment-coded, 3-level factor) as fixed effects. We additionally fit a random intercept for participant and model to account for individual variability.

For each domain (language and vision), we fit two models:

1. Similarity-only model: Included only the median inter-stimulus similarity as a fixed effect.
2. Combined model (Set + Similarity): Included both set type and median inter-stimulus similarity as fixed effects. This model assessed the unique explanatory contribution of set and evaluated whether set explained brain predictivity over and above stimulus similarity.

Each LME model was fit using the *lme4* package in R and estimated using maximum likelihood. The Pearson correlation values from the encoding analyses were predicted across all language (or vision) ANN models and participants. In Experiment 2, this yielded 252 conditions (12 subjects × 7 models × 3 set types), and in Experiment 3, 84 conditions (4 subjects × 7 models × 3 set types). We compared model fits using the likelihood ratio test (with the *anova* function from the *lme4* package in R).

## 4. Behavioral data collection

### Experiment 2, behavioral data collection and analysis

We measured how humans perceived sentences in Experiment 2 (high-agreement, random, and low-agreement) by collecting a set of behavioral ratings. Ratings were collected for the following 10 variables:

We recruited participants through the Prolific platform (www.prolific.com). The full set of 240 sentences in Experiment 2 (80 sentences in each of the three sets: high-agreement, random, and low-agreement) was divided into two groups of 120 sentences (40 sentences from each condition), and the sentence order was randomized as they were presented to participants. Participants rated each sentence on a scale from 1 (low) to 7 (high) based on a given prompt question. We recruited participants who self-reported English as their first language and had completed at least 100 prior submissions with a hit rate of 95% or higher. At the end of the experiment, participants completed six prompt sentences to assess their language ability and attention levels. A CAPTCHA was included at the beginning of the experiment to prevent bots.

The studies were conducted with approval from and in accordance with MIT’s Committee on the Use of Humans as Experimental Subjects (protocol number 2010000243). The participants gave informed consent before starting each experiment and were compensated for their time (minimum US$12 per hour) (Tuckute *et al*., 2024).

To ensure the quality of the behavioral responses, we inspected several condition-independent metrics. For each experiment (corresponding to one of the rating dimensions above), we inspected the following measures: 1. the pattern of responses over trials, 2. the response distribution, 3. the autocorrelation between temporally adjacent responses, 4. the distribution of reaction times, 5. the correlation of individual ratings with the average ratings of other participants, and 6. responses to the sentence completion task. We excluded participants if: a. their sentence completions were not fluent in English or appeared to be generated by language models; b. their pattern of responses was flat or their response distribution substantially differed from the population-level distribution; c) their reaction times were unusually long compared to the population; or d. their responses showed a low correlation with the population average. This procedure led to the exclusion of 95 out of 360 participants.

We then computed the ratings for all sentences for each behavioral metric. To combine the ratings across participants, we z-scored the responses for each participant across all sentences.

Next, we calculated the average rating for each sentence set (high-agreement, random, and low-agreement) for each participant and combined these averages to obtain a population-level rating.

### Experiment 3, behavioral data collection and analysis

We measured how humans perceived images by collecting a set of behavioral ratings. Ratings were collected for the following 10 variables:

| Behavioral metric | Question |
| --- | --- |
| Composition<br>[From | How good is the composition of the image? |
| Semantic coherence | How likely are the different components of the image to appear together? |
| Mental state content | How much does the image make you think of other people's experiences, thoughts, beliefs, desires, and/or emotions? |
| Physical world content | How much does the image make you think of physical objects and/or physical causal interactions? |
| Arousal | How much does the image make you feel stimulated, excited, frenzied, wide-awake, and/or aroused? |
| Valence | How much does the image make you feel happy, pleased, content, and/or hopeful? |
| Frequency | How much does the image “make sense”, depicting things and events that you are likely to encounter in the real world? |
| Expansiveness | How zoomed-out vs. zoomed-in is this image? |
| Clutter | How visually cluttered is the image? |
| Naturalness | How natural (not human-made) are the objects/events that the image depicts? |
| Aesthetics | How aesthetically pleasing is the image? |
| Typicality | How typical is the scene in the image? |

We recruited participants through the Prolific platform (www.prolific.com). In addition to the 240 images, we repeated 20 images from the random set to assess the consistency of participants’ ratings. Participants rated each image on a scale from 1 (low) to 7 (high) based on a given prompt question. We recruited participants who self-reported English as their first language, had a hit rate of 95% or higher in their prior submissions, and had completed at least 100 prior submissions. At the end of the experiment, participants were asked to report whether they observed any of the eight prompt images (four of which were in the main experiment, and four were not) to assess their level of attention. A CAPTCHA was included at the beginning of the experiment to ensure that no bots participated. We collected behavioral ratings from a pool of n = 243 participants.

The studies were conducted with approval from and in accordance with MIT’s Committee on the Use of Humans as Experimental Subjects (protocol number 2010000243). The participants gave informed consent before starting each experiment and were compensated for their time (minimum US$12 per hour) (Tuckute *et al*., 2024).

To ensure the quality of behavioral responses, we inspected a number of condition independent metrics to judge participants’ level of attention. For each experiment, we evaluated the following measures: 1. the pattern of responses over trials, 2. the response distribution, 3. the correlation between the two repetitions of the 20 random images, 4. the distribution of response times, and accuracy in identifying images that were used in the experiment. We adopted a liberal approach to excluding participants, considering a participant as an outlier only in the following cases: a. if their rating consistency for repeated images was not significantly different from random, b. if their pattern of responses was flat, or c. if their performance in identifying images was not different from chance. Based on these criteria, we excluded 29 participants from the pool of 243 participants.

We then computed ratings for images based on each behavioral metric. To compare and pool responses across participants, we z-scored the responses for each participant across all image conditions. Next, we calculated the average rating for each condition for each participant and combined these averages to obtain a population-level rating.

## 5. Statistical testing

As noted in the main text, we primarily relied on Mann-Whitney U tests to compare the distributions of values between conditions, stimulus sets, and models’ predictivity values. Below, we outline the strategies used to perform statistical comparisons with the Mann-Whitney U test in cases where direct comparisons between conditions were not feasible.

### Experiment 1: Comparing linguistic feature values between the random and low-agreement sentences

To assess whether the distribution of linguistic features in the low-agreement set differed from what would be expected under random sampling, we implemented a non-parametric procedure using the Wilcoxon rank-sum test. First, we computed the rank-sum statistic between feature values in low-agreement sentence set and those in the full corpus. Next, we generated a null distribution of rank-sum statistics by drawing 200 random sentence sets from the corpus, each with the same number of sentences as the low-agreement set. For each of these random sets, we calculated the rank-sum statistic against the full corpus. Finally, we calculated the proportion of random sentence sets with rank-sum statistics greater than or equal to the observed rank-sum statistic for low-agreement sentence set. This proportion served as our empirical p-value to infer whether the observed linguistic values in a low-agreement set is expected by chance.

### Experiment 1: Comparing the distributions of RDM values between the random and low-agreement sentences

To assess whether the distribution of sentence dissimilarities in the RDM of the low-agreement set differed from what would be expected under random sampling, we implemented a nonparametric procedure using the Wilcoxon rank-sum test. For each language model, we first constructed the full-set RDM by extracting the upper-triangular part of the RDM for all 8,409 sentences (35,351,436 values) and subsampled 1 million values at random for computational efficiency. We also created a group of random-set RDM values by drawing 200 random subsets of 200 sentences each from the full set and similarly extracting their upper-triangular parts (19,900 values per set). We then computed the rank-sum statistic between the upper-triangular RDM values of the low-agreement set (19,900 values) and the full-set RDM values. To generate a null distribution, we calculated the rank-sum statistic between each of the 200 random-set RDMs and the full-set RDM values. Finally, we calculated the proportion of random sentence sets with rank-sum statistics greater than or equal to the observed rank-sum statistic for the low-agreement sentence set. This proportion served as our empirical p-value, allowing us to infer whether the observed RDM values in the low-agreement set were expected by chance.

### Experiment 2: Comparing linguistic feature values between the different set of sentences

Here we employed 2 methods to compare high-agreement, random, and low-agreement sentences. First, we compared each set with other sets using Mann-Whitney U test. Second, to assess whether the distribution of linguistic features in each set differed from what would be expected under random sampling, we implemented a non-parametric procedure using the Wilcoxon rank-sum test. First, we computed the rank-sum statistic between feature values in each sentence set and those in the full corpus. Next, we generated a null distribution of rank-sum statistics by drawing 200 random sentence sets from the corpus, each with the same number of sentences as the optimized set. For each of these random sets, we calculated the rank-sum statistic against the full corpus. Finally, we calculated the proportion of random sentence sets with rank-sum statistics greater than or equal to the observed rank-sum statistic for the set. This proportion served as our empirical p-value to infer whether the observed linguistic values in high-agreement, random, and low-agreement sets are expected by chance.

### Experiment 2: Comparing the distributions of RDM values of the high-agreement, random, and low-agreement sentences with RDM values from random samples of sentences

To assess whether the distribution of sentence dissimilarities in the RDMs of the high-agreement, random, and low-agreement sets differed from what would be expected under random sampling, we implemented a nonparametric procedure using the Wilcoxon rank-sum test. For each language model, we first constructed the full-set RDM by extracting the upper-triangular part of the RDM for all 5,319 sentences (14,137,903 values) and subsampled 1 million values at random for computational efficiency. We also created a group of random-set RDM values by drawing 200 random subsets of 80 sentences each from the full set and similarly extracting their upper-triangular parts (3,160 values per set). We then computed the rank-sum statistic between the upper-triangular RDM values of each set (high-agreement, random, and low-agreement; 3,160 values per set) and the full-set RDM values. To generate a null distribution, we calculated the rank-sum statistic between each of the 200 random-set RDMs and the full-set RDM values. Finally, we calculated the proportion of random sentence sets with rank-sum statistics greater than or equal to the observed rank-sum statistic for each set. This proportion served as our empirical p-value, allowing us to infer whether the observed RDM values in each set were expected by chance.

### Experiment 3: Comparing the distributions of RDM values of the high-agreement, random, and low-agreement images with RDM values from random samples of images

To assess whether the distribution of image dissimilarities in the RDMs of the high-agreement, random, and low-agreement image sets differed from what would be expected under random sampling, we implemented a nonparametric procedure using the Wilcoxon rank-sum test. For each model, we first constructed the full-set RDM by extracting the upper-triangular part of the RDM for all 1,000 NSD images (499,500 values) and subsampled 1 million values at random for computational efficiency. We also created a group of random-set RDM values by drawing 200 random subsets of 80 images each from the full set and similarly extracting their upper-triangular parts (3,160 values per set). We then computed the rank-sum statistic between the upper-triangular RDM values of each set (high-agreement, random, and low-agreement; 3,160 values per set) and the full-set RDM values. To generate a null distribution, we calculated the rank-sum statistic between each of the 200 random-set RDMs and the full-set RDM values.

Finally, we calculated the proportion of random image sets with rank-sum statistics greater than or equal to the observed rank-sum statistic for each set. This proportion served as our empirical p-value, allowing us to infer whether the observed RDM values in each set were expected by chance.

## Supplementary figures

**Supplementary Figure 1.**
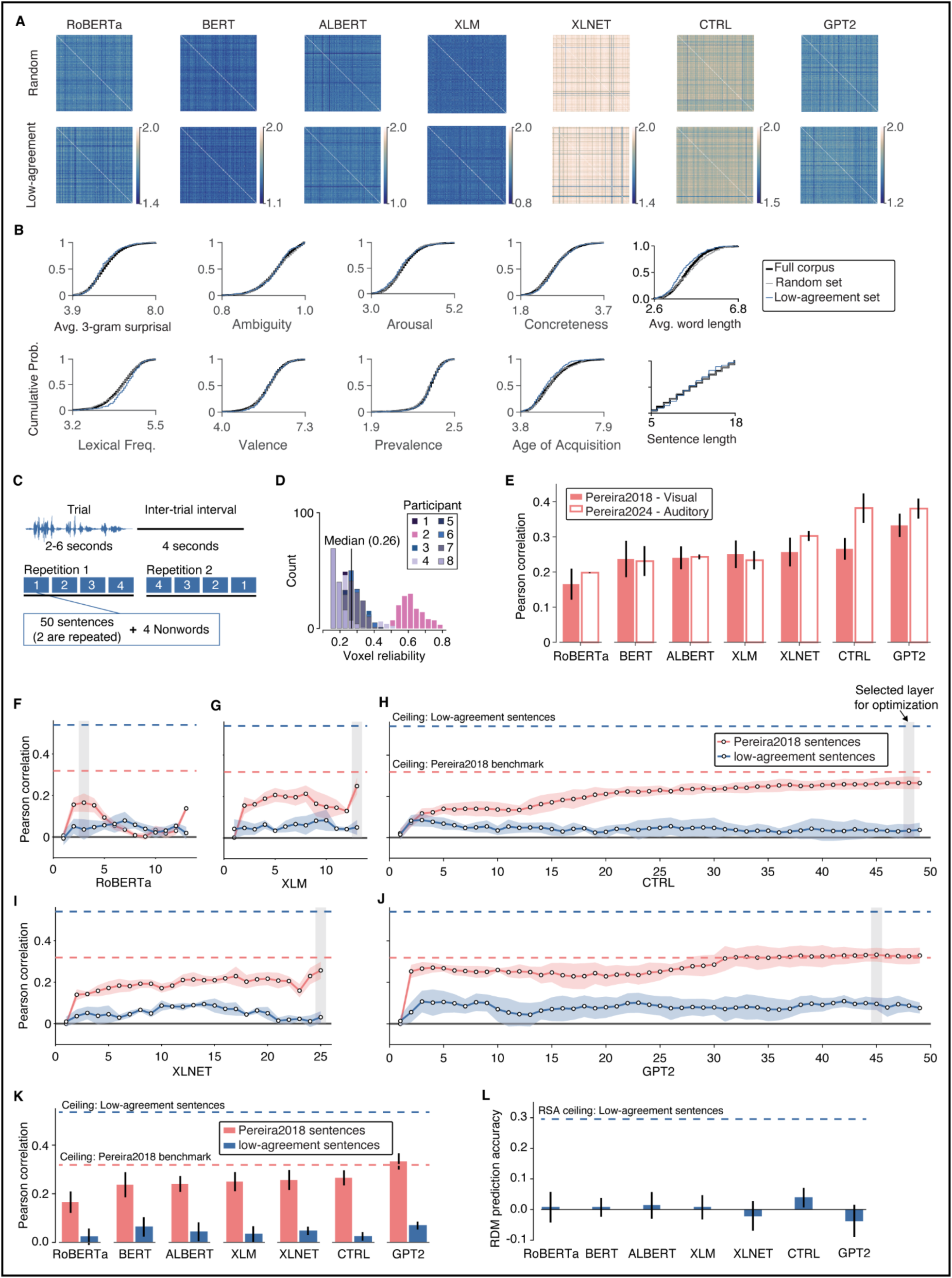
**A.** Sentence RDMs in the random and low-agreement sets for each model. For most models, the distribution of RDM values for the low-agreement set was similar to what would be expected from a random sample (empirical p-values: RoBERTa=0.045; BERT=0.01; ALBERT=1.0; XLM=0.96; XLNET=0.36; CTRL=0.45; GPT2=0.2). **B.** Empirical cumulative probability distributions for several lexical features in the full corpus, the random set, and the low-agreement set (empirical p-values for random vs. low-agreement: avg. 3-gram surprisal=0.93; ambiguity=0.6; arousal=0.98; concreteness=0.09; avg. word length=1.0; lexical frequency<0.001; valence=0.48; prevalence=0.33; age of acquisition=0.99; sentence length=0.73). No significant differences were observed between the random and low-agreement sets, except for lexical frequency. **C.** Experimental design in Experiment 1. Each trial lasted between 2 and 8 seconds, followed by an inter-trial interval of 4 seconds. Each session consisted of 2 repetitions of 200 sentences and 16 nonword sequences, distributed across 8 runs. The order of stimuli in each run was fixed, and the runs were presented in palindromic order [1,2,3,4,4,3,2,1]. **D**. Distribution of voxel reliability across the two repetitions of sentences in each of the 8 participants. **E.** Consistency in voxel predictivity between visual presentations of sentences. In Pereira 2024, we collected voxel responses from a sample of 192 sentences that were used in experiment 2 of Pereira 2018 dataset. Sentences selected such that they covered all passages and topics from the original experiment. Each auditory sentence was 4 seconds long, and model predictivities in the Pereira2024-Auditory experiment were comparable to those in the Pereira2018-Visual experiment. **F.** Layerwise voxel predictivity for the Pereira2018 sentences and the low-agreement sets in the RoBERTa model, showing that the layer used for stimulus optimization is shaded in gray **G–J.** Same as (F) but for XLM, CTRL, XLNET, and GPT2, respectively. **K, L.** Models’ ability to predict voxel activation under an alternative voxel reliability criterion. Instead of selecting the top 10% of voxels by reliability, we used a threshold of 0.1. The pattern replicates the findings shown in Figure 2.

**Supplementary Figure 2.**
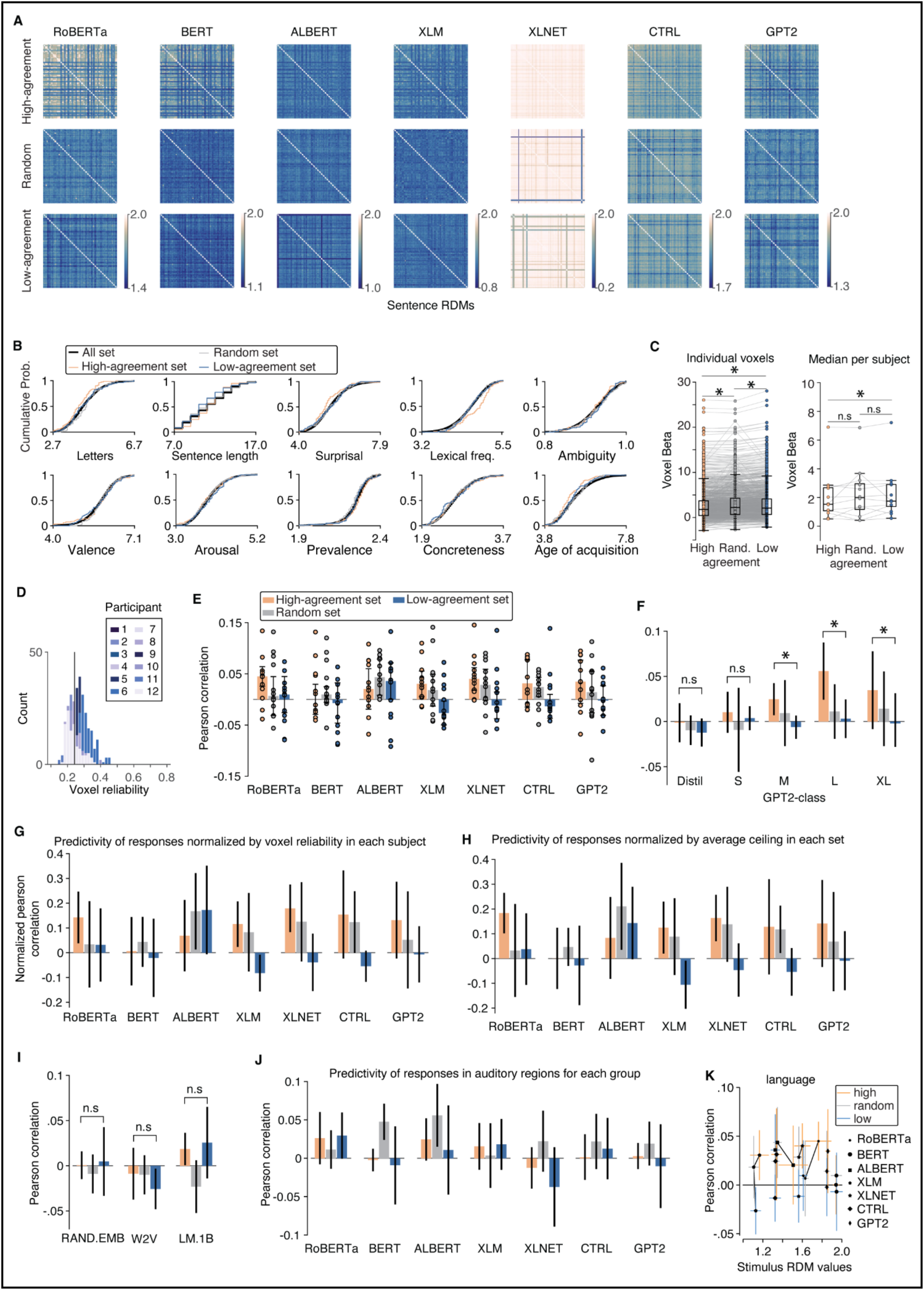
**A.** Sentence RDMs in high-agreement, random, and low-agreement sets for each model. For all models, the distribution of RDM values for the low-agreement set was similar to what would be expected from a random sampling (empirical p-values for comparison between each set and random sampling: RoBERTa: [high=1.00, random=0.88, low=0.09]; BERT: [high=1.00, random=0.90, low=0.08]; ALBERT: [high=0.87, random=0.48, low=0.14]; XLM: [high=1.00, random=0.80, low=1.00]; XLNET: [high=0.12, random=0.07, low=0.15]; CTRL: [high=0.96, random=0.29, low=0.40]; GPT2: [high=0.99, random=0.50, low=0.38]). **B.** Empirical cumulative probability distributions for several lexical features in the full corpus, the random set, and the low-agreement set. The p-values listed are for comparisons between each set and random sampling: avg. 3-gram surprisal: [high=0.99, random=0.45, low=0.55]; ambiguity: [high=0.73, random=0.22, low=0.18]; arousal: [high=0.73, random=0.70, low=0.14]; concreteness: [high=0.55, random=0.74, low=0.79]; avg. word length: [high=1.00, random=0.10, low=0.70]; lexical frequency: [high<0.001, random=0.51, low=0.38]; valence: [high=0.21, random=0.35, low=0.48]; prevalence: [high=0.04, random=0.96, low=0.57]; age of acquisition: [high=1.00, random=0.57, low=0.85]; sentence length: [high=0.44, random=0.84, low=0.95]. No significant differences were observed between the random set and each experimental set, except for lexical frequency and prevalence in the high-agreement set. **C.** Voxel activation patterns in language-responsive regions for high, random, and low-agreement sentences, shown across voxels (left) and participants (right). The median/standard deviation of activation across all 1086 voxels is [high=1.77/3.60, random=2.21/3.73, low=2.11/3.76]; across all 12 participants, [high=1.52/1.66, random=1.99/1.72, low=1.74/1.69]. **D**. Distribution of voxel reliability across the two repetitions of the random set of sentences in each of the 12 participants. **E.** Performance of ANN language models in predicting voxel responses in language-responsive regions for the high-agreement (orange) and low-agreement (blue) sentences, with performance for the random set shown in grey (same as Figure 3I). Each dot represents an individual participant. **F.** Performance of models not used for stimulus selection in predicting voxel responses in language-responsive regions for each set, including GPT2-class models (Distill, S, M, and L, with XL shown for comparison). Only larger models (M, L) reliably distinguish between high-and low-agreement, while smaller models (Distill, S) do not. **G,** Performance of ANN language models in predicting voxel responses for high-agreement (orange) and low-agreement (blue) sentences, with random set (grey), after normalizing by participant-level ceilings (see Methods). For each repetition, we normalized by each participant’s ceiling, averaged the two repetitions, then took the median/median absolute deviation over all 12 participants (similar to Figure 3I). **H**. Same as G, but here we first computed an average ceiling over the two repetitions, then normalized each participant’s average predictivity by that single ceiling value. The pattern is similar to the main figure results. **I.** Performance of baseline models (random-embedding, word2vec, and an LSTM-based LM1B) in predicting voxel responses in language-responsive regions for the high-agreement (orange) and low-agreement (blue) sentences, with random set performance in grey. **J.** Performance of ANN language models in predicting voxel responses in auditory regions for high-agreement (orange) and low-agreement (blue) sentences, with random set in grey. No difference between high-and low-agreement sets was observed in auditory regions. **K.** Relationship between the distribution of sentence dissimilarities in each model and the prediction of voxel responses for high-agreement, random, and low-agreement sentences.

**Supplementary Figure 3.**
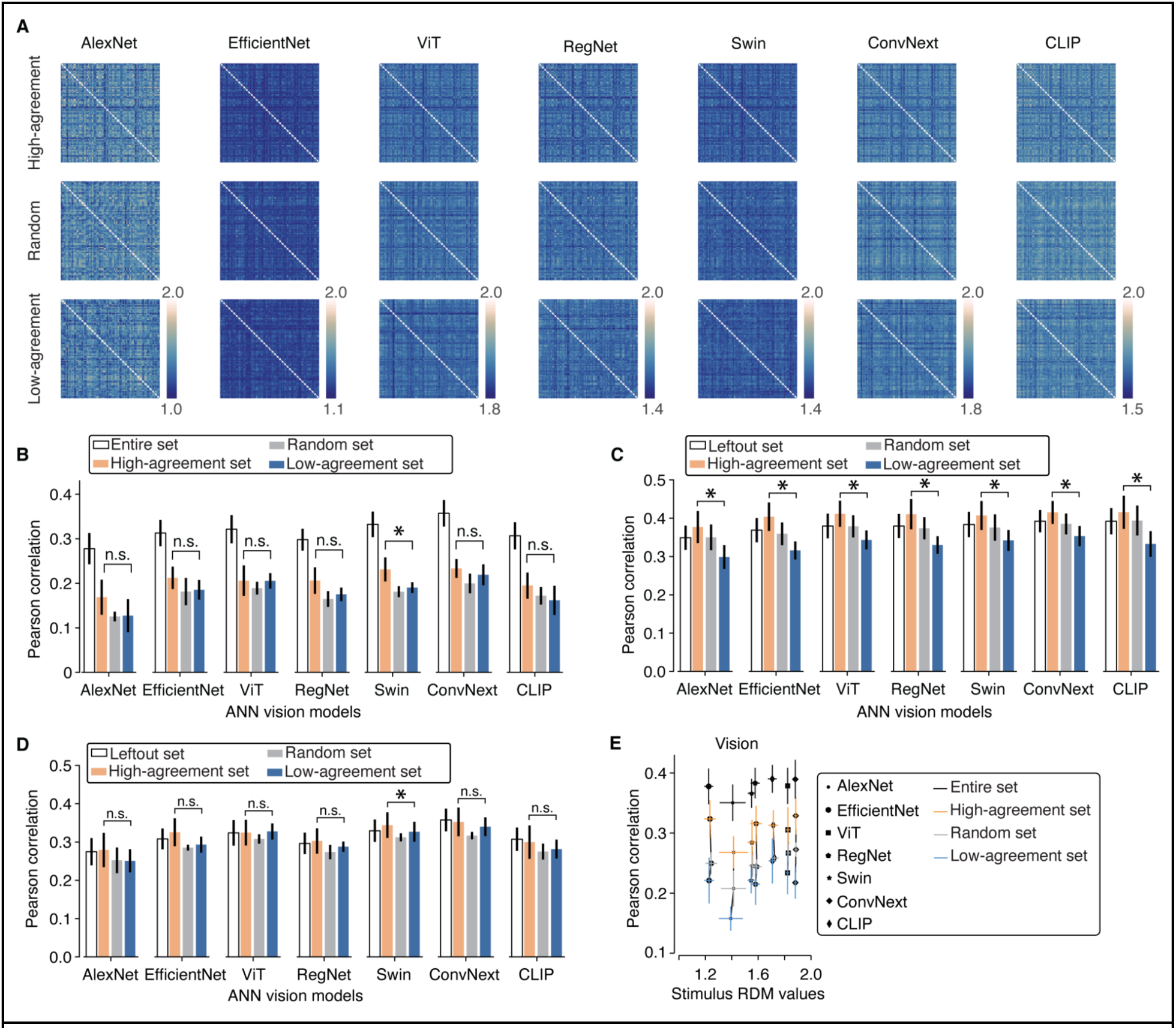
**A.** Image RDMs in high-agreement, random, and low-agreement sets for each model. For all models, the distribution of RDM values for the low-agreement set was similar to what would be expected from a random sampling (empirical p-values for comparison between each set and random sampling: AlexNet: [high=0.82, random=0.88, low=0.29]; EfficientNet: [high=0.77, random=0.99, low=0.50]; ViT: [high=0.67, random=0.97, low=0.36]; RegNet: [high=0.68, random=0.94, low=0.57]; Swin: [high=0.73, random=0.94, low=0.35]; ConvNext: [high=0.84, random=0.96, low=0.39]; CLIP: [high=0.61, random=0.99, low=0.30]). **B.** Performance of ANN vision models in predicting voxel responses in early visual cortex (EVC) for high-agreement (orange) and low-agreement (blue) images, with random-set performance shown in grey (same as Figure 3I). Only one model (Swin) shows a reliable difference between high and low-agreement sets.**C.** Performance of ANN vision models in predicting voxel responses in OTC for high-agreement, random, and low-agreement image sets, using a regression built on a left-out set of 774 images. The regression was trained only on images not included in any of these sets, then tested on the three sets. Consistent with the main figure, OTC responses are best predicted for high-agreement images, followed by random and low-agreement images. **D.** Same analysis as C but for voxels in EVC. Here, there was no reliable relationship between model agreement and voxel predictivity. For six out of seven models, the predictivity for high-agreement images is similar to that for low-agreement images. **E.** Relationship between each model’s distribution of RDM values and its ability to predict voxel responses in OTC. In the visual domain, stimulus optimization was modified to control for shifts in the distribution of RDM values between random and high-or low-agreement images, by adding a penalty based on the KL divergence from an average random-set distribution.

**Supplementary figure 4.**
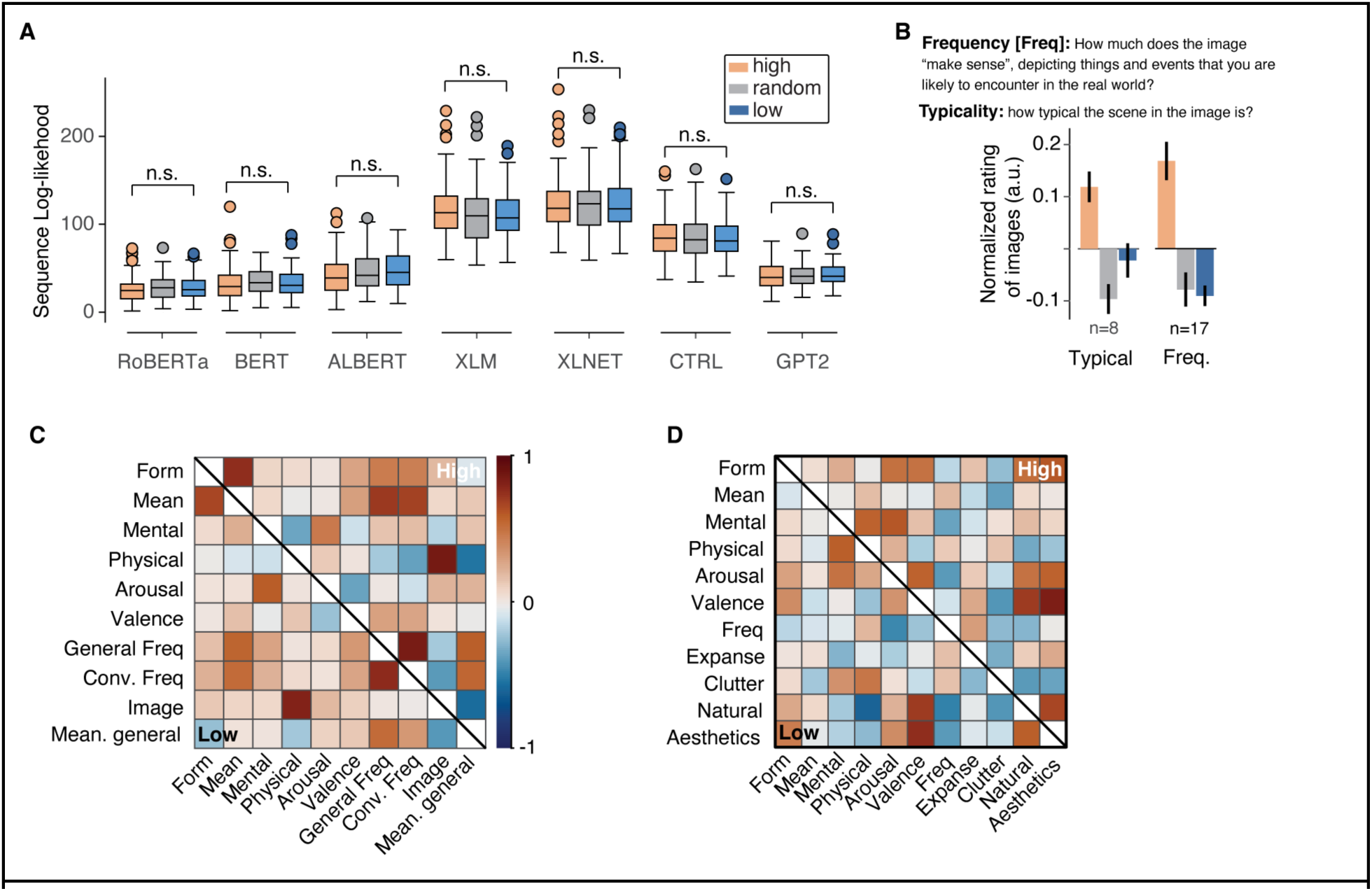
**A.** Model behavior for different sets of sentences. We computed model sequence log-likelihoods for all sentence sets and found that the models did not differentiate between high-and low-agreement sentences; both sets yielded similar likelihood values. This suggests that, from the models’ perspective, there is no clear behavioral distinction between high-and low-agreement sentences. **B.** Behavioral rating of images along frequency and typicality. Neither of these patterns replicated the results observed for the language domain. **C.** Covariation across dimensions in the language domain. Even though General Frequency and Meaning Generality are moderately correlated, their correlations with other factors differ, suggesting they each explain unique variance in the stimulus sets. **D.** Same as C but for dimensions in the visual domain.

**Supplementary Figure 5.**
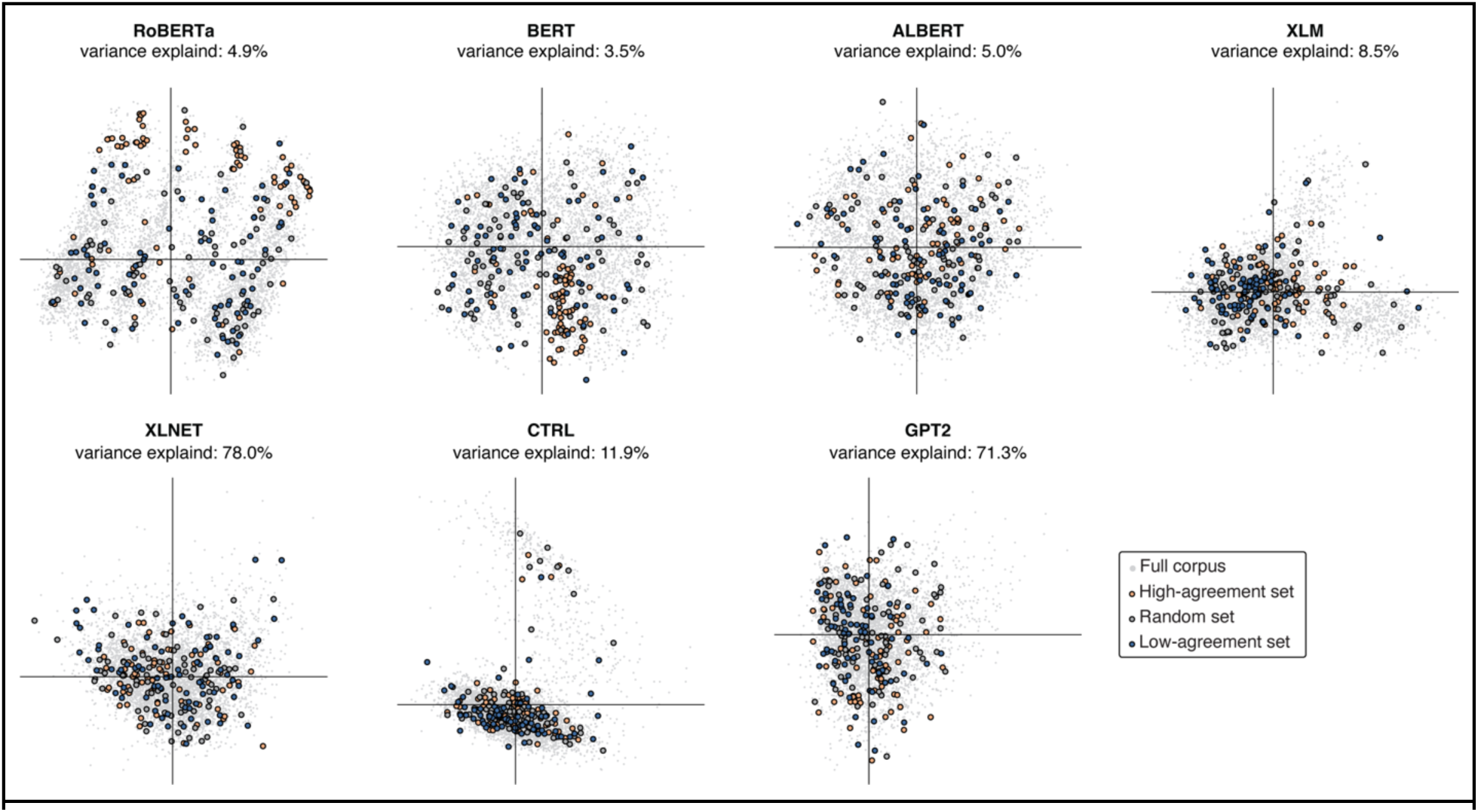
Organization of the stimulus set in the representation of language models (Experiment 2). For each of the seven language models, we performed a principal component analysis (PCA) on the representations of 5,319 sentences (excluding those in the high-agreement, random, and low-agreement sets). We then used the resulting principal axes to project sentences from each set onto this low-dimensional embedding. The first two principal components, along with their explained variance, are shown here. In many of the models, all sentence sets span the entirety of this two-dimensional space, with no clear boundary separating the sets—indicating that neither the high-nor the low-agreement sentences are outliers in the models’ representations.

